# Coordinated subpocket engagement underlies nitazene potency at the µ-opioid receptor

**DOI:** 10.64898/2026.04.27.721072

**Authors:** Margaret Jane Robinson, Li Chen, Abhishek Thakur, Kuo-Hao Lee, Lei Shi

**Author notes:** equal contribution (Margaret Jane Robinson, Li Chen, Abhishek Thakur).

## Abstract

Nitazenes are emerging synthetic opioids that exhibit exceptionally high potency at the µ-opioid receptor (MOR) and contribute to rising overdose fatalities worldwide. Despite extensive *in vitro* profiling, the structural determinants underlying their structure–activity relationships (SARs) remain unresolved. Here, we combine functional profiling with quantum mechanical calculations and molecular dynamics (MD) simulations to establish a MOR structure-based SAR for nitazenes. Across functional assays, systematic variation at the R1, R2, and R3 positions revealed non-additive effects on potency and identified optimal R1 chain length, R2 N-desethylation, and retention of the 5-nitro group as key determinants of high MOR potency. Consistent with this framework, N-desethyl isotonitazene emerged as the most potent analogue. Structural analysis of the cryo-EM MOR–G_i_–fluornitrazene complex, together with MD simulations of multiple nitazene analogues, revealed a conserved trivalent binding architecture in which each substituent engages distinct subpockets. N-desethylation at R2 increases the positive electrostatic surface at the protonated amine, reduces steric constraints near transmembrane helix (TM) 7, strengthens R3 interactions, and allosterically modulates R1 engagement in a substituent-dependent manner. Additionally, optimal R1 chain length and shape stabilize the TM5–TM6 interface and influence activation-relevant TM6 dynamics, defining a unified SAR at R1 across nitazene and fentanyl scaffolds. Together, these findings indicate that nitazene potency reflects substituent-dependent coupling among R1, R2, and R3 within the MOR binding pocket, with R3 engagement distinguishing nitazenes from fentanyl. This framework establishes a coherent structural model of nitazene–MOR recognition that accounts for their unusually high potency and efficacy.

## INTRODUCTION

The “opioid epidemic” is a public health crisis characterized by widespread misuse of opioids and rising overdose deaths.^1^ The epidemic’s third wave, which began around 2013 with the emergence of illicitly manufactured fentanyl and its analogues, has continued to intensify into the 2020s.^1, 2^ While fentanyl is used clinically as a potent analgesic, it has high abuse liability that arises from activation of the μ-opioid receptor (MOR). MOR activation produces profound analgesia compared to other opioid receptors but is also primarily responsible for adverse effects of opioids, including reward-seeking behaviors and, most dangerously, life-threatening respiratory depression.^3–5^

Fentanyl’s widespread use and subsequent proliferation of related analogues prompted swift international scheduling over the past decade, spurring the emergence of structurally diverse novel synthetic opioids (NSOs) designed to evade regulatory control.^6, 7^ Among them, the appearance of isotonitazene (INZ) in illicit markets in 2019 marked the reemergence of the nitazenes (2-benzylbenzimidazole derivatives).^8, 9^ In the years since, nitazenes have become a rapidly growing global NSO threat, with increasing involvement in opioid-related fatalities reported across multiple countries.^10^ Like fentanyl, nitazenes are highly potent MOR agonists, with several analogues surpassing fentanyl in potency, efficacy, and lipophilicity, thereby exacerbating MOR-mediated adverse effects and increasing abuse liability.^11–13^ Their rapid diversification has prompted extensive side-by-side *in vitro* profiling of nitazenes and fentanyl analogues to understand their differences;^13^ moreover, recent efforts also profiled “prophetic” nitazenes predicted to emerge imminently in illicit markets.^14^

Among nitazene analogues, N-desethyl isotonitazene (DINZ) is of particular concern. Although DINZ is a metabolite of INZ, *in vitro* analyses revealed that DINZ exhibits greater potency than most of the other nitazenes.^13, 15^ Early drug surveillance programs in the United States have reported increasing detection of DINZ in drug samples and overdose cases in the absence of INZ.^16^ Notably, between October 2023 and April 2024 alone, National Medical Service Labs reported DINZ in 24 post-mortem cases without concurrent detection of INZ,^17^ underscoring the independent threat posed by DINZ as an abused NSO.

The consensus from *in vitro* and *in vivo* studies supports a consistent structure–activity relationship (SAR) defined by substituents at three key positions on the nitazene scaffold (Figure 1): the alkoxy tail (R1) on the 2-benzyl substituent of the benzimidazole core, the protonated amine and its alkyl substitutions (R2), and the 5-nitro group (R3) on the benzimidazole core. Functional potency decreases as the length of R1 deviates from the optimal isopropoxy or ethoxy groups.^18^ The alkyl substitutions at R2 can potentially modulate the orientation of the protonated amine, which mediates the key ionic interaction with Asp149^3.32^ of MOR (superscripts denote Ballesteros−Weinstein numbering^19^). Interestingly, whereas INZ is less potent than DINZ, etonitazene (ENZ) is more potent than its N-desethyl analogue (DENZ),^11, 13, 20^ suggesting that R1 and R2 are allosterically coupled. In addition, an early study revealed the importance of the 5-nitro group (R3) on the benzimidazole core in promoting antinociceptive activity,^21^ and recent *in vitro* studies confirmed that removal of the 5-nitro group reduces MOR activation.^15, 22, 23^

**Figure 1:**
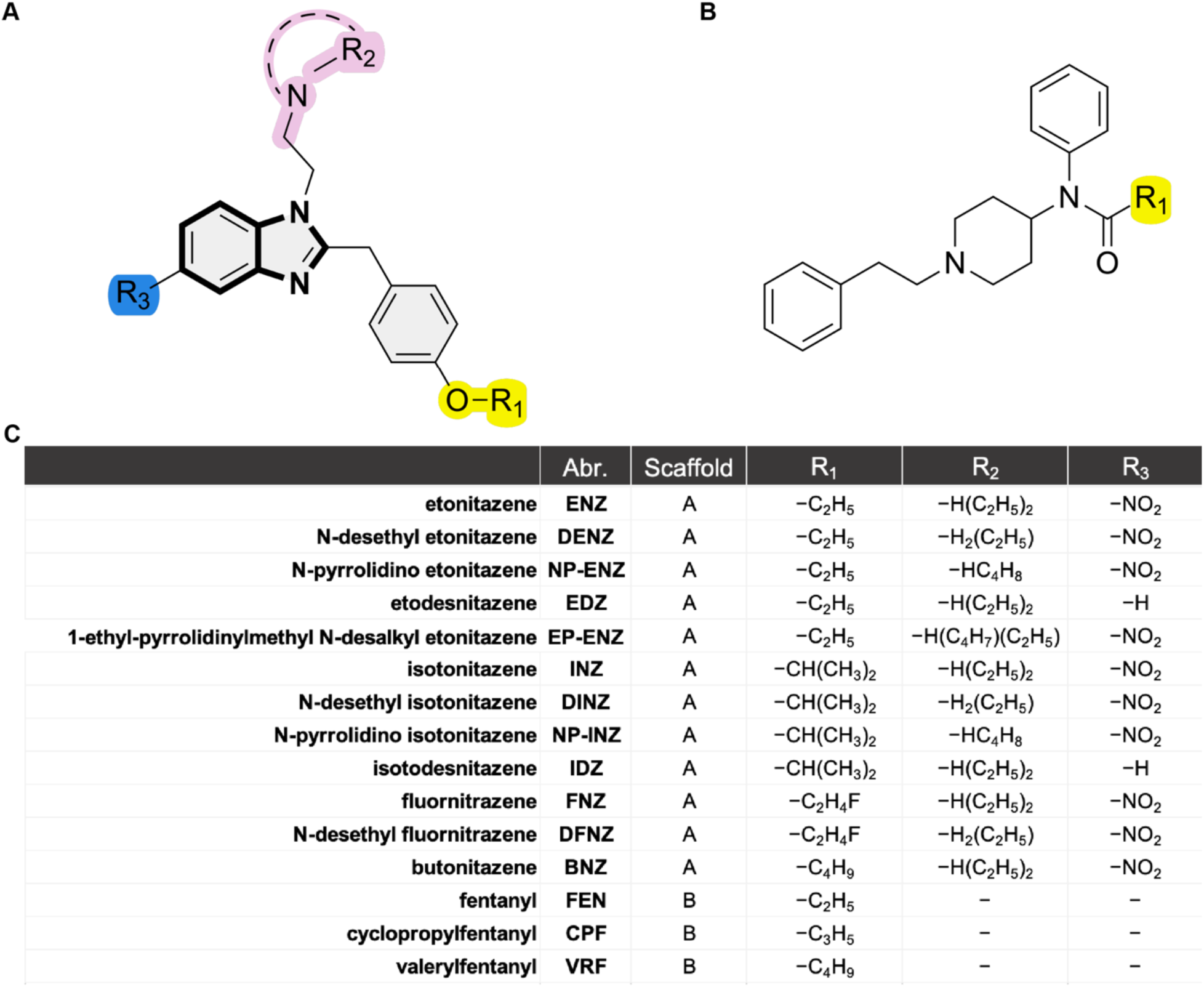
Chemical structures of nitazenes and fentanyls investigated in this study. Analogues were systematically selected to explore variations of the parent scaffolds, shown for nitazenes in (A) and fentanyls in (B). Site-specific modifications are shown in (C). Full chemical structures for all tested compounds are provided in Figure S1. In the nitazene scaffold shown in (A), the yellow, pink, and blue regions, together with the bolded atoms, define the R1, R2, R3, and core regions used for regional contact frequency analysis in our modeling work (see Figures 5, 6, and S5). Note that the aromatic ring of the 2-benzyl substituent is not included in any region but is included in the whole-molecule contact frequency calculations shown in Table 2.

The recently reported cryo-EM structure of the MOR–G_i_–fluornitrazene (FNZ) complex^24^ defined the binding mode of FNZ, in which the R1 fluoroethoxyl group on the 2-benzyl substituent and R2 diethylaminoethyl substituent attached to the benzimidazole core occupy the orthosteric binding site (OBS), whereas the R3 5-nitro group and the benzimidazole core itself (from which the nitazene name derives) extend unexpectedly beyond this pocket. In this binding mode, each functional group (R1, R2, and R3; Figure 1) engages distinct subpockets within the binding site. Notably, the R1 of FNZ occupies the same subpocket between the transmembrane helices (TMs) 5 and 6 that accommodates the N-propionamide tail of fentanyl bound to MOR,^25^ suggesting related SAR features between the nitazene and fentanyl scaffolds.

FNZ and N-desethyl fluornitrazene (DFNZ) have demonstrated an improved safety profile, lower liability for abuse, and markedly reduced respiratory depression in animal models.^24^ While ongoing profiling of nitazenes continues to affirm the substantial dangers of this class, the discovery of FNZ and DFNZ complicates how nitazene SAR relates to safety.^24^ This dual reality, therapeutic promise amid rising nitazene-related fatalities, underscores the urgent need for a mechanistic, structure-based understanding of the MOR-nitazene interactions to distinguish therapeutic from adverse effects and to better inform public health and pharmacological strategies.

To address these uncertainties in nitazene pharmacology, we sought to establish a MOR structure-based SAR for the nitazene scaffold. We combined *in vitro* functional assays with *in silico* modeling to relate experimental measures of potency and efficacy to underlying molecular interactions at MOR. Conformational and quantum-mechanical analyses were conducted to characterize how N-desethyl modifications alter electronic and structural properties, and MD simulations were used to assess their potential impact on receptor–ligand interactions. Comparison of the nitazene and fentanyl scaffolds within the MOR binding pocket was undertaken to identify shared structural determinants of activity. This integrated approach establishes a framework for dissecting the molecular features that may help to distinguish harmful from potentially therapeutic opioid actions.

## RESULTS

To evaluate how modifications at the R1, R2, and R3 positions influence nitazene activity, we employed three *in vitro* functional assays at MOR, including two Homogeneous Time-Resolved Fluorescence (HTRF)-based assays (adenosine cyclic 3′,5′-phosphate (cAMP) inhibition and GTP-G_i_ binding) and a Bioluminescence Resonance Energy Transfer (BRET) assay (Gα_oA_(Go) activation), each capturing a distinct G-protein signaling outcome as described in the following sections. In each assay, concentration–response data for every tested opioid were fitted to nonlinear regression models to determine the maximum response (%E_max_, normalized to a reference full agonist [D-Ala^2^,N-MePhe^4^,Gly-ol^5^]enkephalin (DAMGO)) and the concentration producing half-maximal activation (EC_50_), providing estimates of efficacy and potency, respectively.

To systematically elucidate SARs for each R substituent, we selected 12 nitazene analogues with stepwise modifications at the R1, R2, and R3 positions. Using ENZ, INZ, or FNZ as the starting structures, each analogue differs by a single substitution from these canonical nitazenes (see Figures 1 and S1). To compare nitazene activity profiles with those of other common NSOs and clinical analgesics, we also included three fentanyl analogues with structural modifications analogous to the R1 position of nitazenes, and two phenanthrene (morphinean) compounds, morphine and buprenorphine, both established clinical analgesics.^26, 27^

### HTRF cAMP inhibition defined relative potency ranking of nitazenes

In the HTRF-based cAMP inhibition assay, extending our previous analysis^13^ to additional analogues, all 12 nitazenes displayed lower EC_50_ values than the reference full agonist DAMGO (EC_50_ = 1.7 nM) (Figure 2A; Table 1). Among nitazenes, DINZ was the most potent compound, with an EC_50_ of 13 pM, whereas butonitazene (BNZ) was the least potent (EC_50_ = 1.3 nM). The remaining nitazenes clustered into two potency groups divided by a 3-fold difference in EC_50_. isotodesnitazene (IDZ), etodesnitazene (EDZ), and 1-ethyl-pyrrolidinylmethyl N-desalkyl etonitazene (EP-ENZ) clustered near BNZ, with EC_50_ values ranging from 0.42 to 0.74 nM. All other nitazene analogues displayed EC_50_ values between 0.052 and 0.15 nM, approximately 4-fold higher than DINZ. Notably, fentanyl (EC_50_ = 0.42 nM) fell within the less potent cluster, while its analogue cyclopropylfentanyl (CPF) (EC_50_ = 0.15 nM) grouped with the more potent nitazenes. Buprenorphine (EC_50_ = 0.68 nM) showed potency comparable to less potent nitazenes, whereas valerylfentanyl (VRF) and morphine were less potent than DAMGO by approximately 17- and 3-fold, respectively.

**Figure 2:**
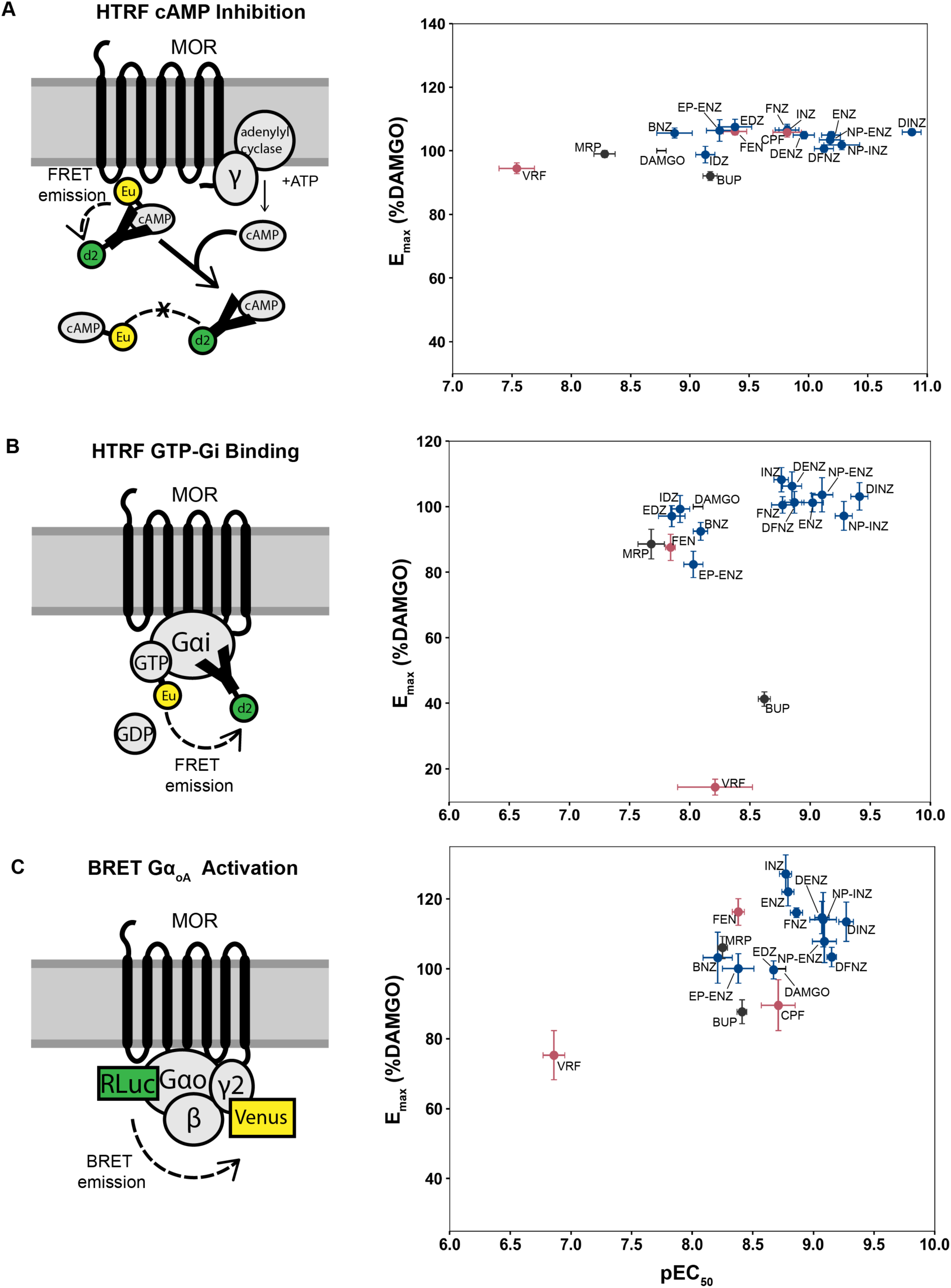
*In vitro* activities of the tested opioids. Concentration-response curves were determined for HTRF cAMP inhibition (A), HTRF GTP-G_i_ binding (B), and BRET Go activation (C). The cartoons depict the luminescence- or fluorescence-labeled constructs used in each assay. The cAMP inhibition cartoon shown in (A) was adapted from “cAMP Guide to optimizing agonists of Gαs” (Revvity, Waltham, MA, USA), and the GTP-G_i_ binding cartoon shown in (B) was adapted from “GTP-Gi binding assay: A guide to optimizing agonists of Gαi” (Revvity, Waltham, MA, USA). In each 2D plot, each point represents one opioid, positioned according to its mean E_max_ (% of DAMGO) and pEC_50_ measured in that assay (see Table 1). Error bars indicate the standard error of the mean (SEM). Nitazenes are shown in blue, fentanyls in red, and reference compounds in black. Note that, across panels, the X-axes use the same span, and the Y-axes use the same span, although the absolute axis limits may differ, highlighting differences in assay resolution for both potency and efficacy.

**Table 1:** E_max_ and EC_50_ values of tested compounds for the indicated assays.

|  | HTRF cAMP Inhibition |  |  |  |  |  | HTRF GTP-Gi Binding |  |  |  |  |  | BRET Go Activation |  |  |  |  |  |
| --- | --- | --- | --- | --- | --- | --- | --- | --- | --- | --- | --- | --- | --- | --- | --- | --- | --- | --- |
|  | pEC50 | SEM | EC50 (nM) | %Emax | SEM | n | pEC50 | SEM | EC50 (nM) | %Emax | SEM | n | pEC50 | SEM | EC50 (nM) | %Emax | SEM | n |
| **DAMGO** | 8.76 | 0.04 | 1.7 | 100.0 | 0.0 | 97 | 8.07 | 0.04 | 8.5 | 100.0 | 0.0 | 47 | 8.73 | 0.04 | 1.9 | 100.0 | 0.0 | 66 |
| **MRP** | 8.28 | 0.09 | 5.2 | 99.0 | 1.1 | 23 | 7.68 | 0.11 | 21 | 88.6 | 4.5 | 6 | 8.25 | 0.04 | 5.6 | 106.1 | 3.2 | 9 |
| **BUP** | 9.17 | 0.06 | 0.68 | 92.1 | 1.4 | 11 | 8.62 | 0.05 | 2.4 | 41.3 | 2.2 | 8 | 8.41 | 0.04 | 3.9 | 87.7 | 3.4 | 11 |
| **ENZ** | 10.19 | 0.08 | 0.065 | 104.9 | 1.1 | 26 | 9.02 | 0.09 | 0.95 | 101.2 | 2.8 | 11 | 8.79 | 0.05 | 1.6 | 122.1 | 4.1 | 14 |
| **DENZ** | 9.96 | 0.09 | 0.11 | 104.9 | 1.2 | 19 | 8.85 | 0.08 | 1.4 | 106.3 | 4.3 | 10 | 9.07 | 0.06 | 0.85 | 114.7 | 4.6 | 13 |
| **NP-ENZ** | 10.18 | 0.09 | 0.066 | 103.4 | 0.9 | 21 | 9.10 | 0.09 | 0.79 | 103.6 | 5.2 | 6 | 9.09 | 0.10 | 0.81 | 107.9 | 6.1 | 10 |
| **EDZ** | 9.38 | 0.14 | 0.42 | 107.5 | 2.4 | 4 | 7.85 | 0.11 | 14 | 97.1 | 3.2 | 7 | 8.67 | 0.01 | 2.1 | 99.7 | 2.6 | 5 |
| **EP-ENZ** | 9.25 | 0.11 | 0.56 | 106.4 | 3.4 | 4 | 8.03 | 0.08 | 9.3 | 82.4 | 4.0 | 7 | 8.38 | 0.13 | 4.2 | 100.1 | 4.2 | 7 |
| **INZ** | 9.82 | 0.07 | 0.15 | 106.5 | 1.3 | 24 | 8.76 | 0.06 | 1.7 | 108.2 | 3.7 | 11 | 8.77 | 0.05 | 1.7 | 127.2 | 5.4 | 13 |
| **DINZ** | 10.87 | 0.08 | 0.013 | 105.9 | 1.0 | 20 | 9.41 | 0.07 | 0.39 | 103.1 | 4.2 | 9 | 9.27 | 0.06 | 0.54 | 113.5 | 5.6 | 16 |
| **NP-INZ** | 10.28 | 0.15 | 0.052 | 101.8 | 1.5 | 5 | 9.28 | 0.07 | 0.52 | 97.2 | 4.4 | 5 | 9.08 | 0.11 | 0.83 | 114.1 | 7.8 | 13 |
| **IDZ** | 9.13 | 0.08 | 0.74 | 98.8 | 2.6 | 4 | 7.92 | 0.08 | 12 | 99.3 | 4.1 | 9 | N.D. | N.D. | N.D. | N.D. | N.D. | N.D. |
| **FNZ** | 9.82 | 0.10 | 0.15 | 106.5 | 1.8 | 6 | 8.77 | 0.09 | 1.7 | 100.5 | 2.5 | 8 | 8.86 | 0.05 | 1.4 | 116.1 | 1.4 | 17 |
| **DFNZ** | 10.13 | 0.08 | 0.074 | 100.7 | 1.2 | 9 | 8.87 | 0.08 | 1.3 | 101.3 | 3.3 | 9 | 9.15 | 0.04 | 0.71 | 103.4 | 2.8 | 16 |
| **BNZ** | 8.87 | 0.15 | 1.3 | 105.6 | 1.6 | 12 | 8.09 | 0.06 | 8.1 | 92.5 | 2.7 | 5 | 8.21 | 0.12 | 6.2 | 103.2 | 7.3 | 8 |
| **FEN** | 9.38 | 0.10 | 0.42 | 106.1 | 1.1 | 19 | 7.84 | 0.04 | 14 | 87.6 | 4.0 | 18 | 8.38 | 0.05 | 4.2 | 116.3 | 3.8 | 17 |
| **CPF** | 9.82 | 0.12 | 0.15 | 105.8 | 1.5 | 12 | N.D. | N.D. | N.D. | N.D. | N.D. | N.D. | 8.71 | 0.14 | 1.9 | 89.6 | 7.3 | 7 |
| **VRF** | 7.54 | 0.15 | 29 | 94.5 | 1.7 | 12 | 8.21 | 0.31 | 6.2 | 14.5 | 2.4 | 4 | 6.86 | 0.09 | 140 | 75.3 | 7.0 | 8 |

N.D., not determined. $E_{\max}$ values are normalized to that of DAMGO. Data are shown as mean ± standard error of the mean
(SEM) for the indicated $n$ replicates.

**Table 2:** Nitazenes consistently engage up to 21 residues in MOR. Contact is defined as any residue with at least one heavy atom within 5 Å of any ligand heavy atom. Only residues with a contact frequency above 0.3 for at least one ligand are included in the table.

|  | ENZ | FNZ | INZ | BNZ | DENZ | DFNZ | DINZ |
| --- | --- | --- | --- | --- | --- | --- | --- |
| <u>Tyr77</u><sup>1.39</sup> | 1.00 | 1.00 | 1.00 | 1.00 | 1.00 | 1.00 | 1.00 |
| <u>Leu123</u><sup>2.57</sup> | 0.96 | 0.99 | 0.95 | 1.00 | 0.99 | 1.00 | 0.99 |
| <u>Gln126</u><sup>2.60</sup> | 1.00 | 1.00 | 1.00 | 1.00 | 1.00 | 1.00 | 1.00 |
| <u>Ser127</u><sup>2.61</sup> | 0.99 | 1.00 | 1.00 | 1.00 | 1.00 | 1.00 | 1.00 |
| <u>Asn129</u><sup>2.63</sup> | 0.14 | 0.51 | 0.01 | 0.01 | 0.02 | 0.00 | 0.00 |
| <u>Tyr130</u><sup>2.64</sup> | 0.99 | 0.99 | 0.99 | 1.00 | 1.00 | 1.00 | 1.00 |
| <u>Leu131</u><sup>2.65</sup> | 0.14 | 0.10 | 0.34 | 0.20 | 0.17 | 0.39 | 0.39 |
| <u>Asp149</u><sup>3.32</sup> | 1.00 | 1.00 | 1.00 | 1.00 | 0.99 | 1.00 | 1.00 |
| <u>Tyr150</u><sup>3.33</sup> | 0.97 | 1.00 | 1.00 | 1.00 | 0.96 | 0.97 | 0.99 |
| <u>Met153</u><sup>3.36</sup> | 0.97 | 1.00 | 1.00 | 1.00 | 0.80 | 0.97 | 0.97 |
| <u>Lys235</u><sup>5.39</sup> | 0.58 | 0.03 | 0.72 | 0.86 | 0.71 | 0.65 | 0.89 |
| <u>Val238</u><sup>5.42</sup> | 0.96 | 0.70 | 0.96 | 0.96 | 0.89 | 0.97 | 0.99 |
| <u>Phe239</u><sup>5.43</sup> | 0.35 | 0.11 | 0.45 | 0.59 | 0.36 | 0.37 | 0.41 |
| <u>Trp295</u><sup>6.48</sup> | 0.33 | 0.57 | 0.76 | 0.76 | 0.20 | 0.72 | 0.42 |
| <u>Ile298</u><sup>6.51</sup> | 0.99 | 1.00 | 1.00 | 0.99 | 0.98 | 0.93 | 0.99 |
| <u>His299</u><sup>6.52</sup> | 0.99 | 0.99 | 0.99 | 0.99 | 0.99 | 0.91 | 1.00 |
| <u>Val302</u><sup>6.55</sup> | 0.97 | 0.93 | 0.93 | 0.88 | 0.92 | 0.78 | 0.97 |
| <u>Trp320</u><sup>7.35</sup> | 0.95 | 0.97 | 0.93 | 0.93 | 0.95 | 0.86 | 0.95 |
| <u>His321</u><sup>7.36</sup> | 0.99 | 1.00 | 0.99 | 1.00 | 0.97 | 0.95 | 0.99 |
| <u>Ile324</u><sup>7.39</sup> | 1.00 | 1.00 | 1.00 | 1.00 | 1.00 | 1.00 | 1.00 |
| <u>Tyr328</u><sup>7.43</sup> | 1.00 | 1.00 | 1.00 | 1.00 | 0.99 | 1.00 | 1.00 |

Underlined residues are within 5 Å of FNZ in the published cryo-EM structure (PDB:9O36)<sup>24</sup>
(see Figure 3).

All nitazenes displayed efficacies comparable to or slightly higher than that of DAMGO, with E_max_ values ranging from 98.8% (IDZ) to 107.5% (EDZ). Both fentanyl (E_max_ = 106.1%) and CPF (E_max_ = 105.8%) showed efficacies comparable to the nitazenes. VRF and buprenorphine had the lowest efficacies (94.5% and 92.1%, respectively). As both VRF and buprenorphine are known partial agonists at MOR,^13, 28–30^ these notably elevated E_max_ values, all above 90%, reflect signal amplification in this assay. Indeed, because cAMP is a secondary messenger (Figure 2A), receptor reserve can amplify the signaling output of this assay, allowing the system to reach maximum response even when only a fraction of receptors is occupied by ligand,^31^ thereby complicating experimental interpretation.^32^

Despite this limitation, the HTRF-based cAMP inhibition assay provides a high signal-to-noise ratio,^13^ effectively enhancing the resolution of potency differences among closely related analogues. This sensitivity yielded a broader and more differentiated distribution of potencies across the tested opioids compared to the other assays used in this study (see below). Accordingly, the relative potency ranking derived from this assay served as the primary reference for the subsequent molecular modeling work to establish the structure-based SAR at MOR described below.

### Low-amplification GTP-G_i_ binding highlighted high intrinsic efficacy of nitazenes

To obtain reliable measurements of ligand efficacy, we employed an HTRF-based GTP–G_i_ binding assay at MOR. This assay measures the GTP–GDP exchange by the Gα_i_ subunit bound to MOR, which occurs proximal to receptor activation and is therefore less susceptible to downstream signal amplification, thereby providing a more accurate estimate of intrinsic efficacy. However, characterizing GTP binding at MOR presented distinct analytical challenges. We observed bell-shaped rather than sigmoidal dose-response curves for some but not all opioids (Figure S2), consistent with a previous report.^33^ To address this issue, we developed a protocol that fits either a sigmoidal or bell-shaped model to each dataset based on goodness of fit (see Methods). This approach yielded consistent measures of potency and efficacy across independent experiments, which were unattainable using the sigmoid model alone, and enabled differentiation among partial, full, and super agonists at MOR (Figure S2).

In this GTP binding assay, we found that nitazenes exhibited efficacies comparable to or exceeding that of DAMGO (Figure 2B; Table 1). The exceptions were BNZ and EP-ENZ, which showed reduced efficacies (92.5% and 82.4% of DAMGO, respectively), likely due to their sterically bulky substituents at the R1 and R2 positions, respectively.

The fentanyl scaffold displayed more divergent efficacies compared to the tightly clustered nitazenes. Fentanyl showed an efficacy of 87.6%, whereas VRF displayed the lowest efficacy of all tested compounds at 14.5% (Figure 2B). This low efficacy exhibited by VRF, together with the relatively low efficacy of buprenorphine (%E_max_ = 41.3%), indicates limited signal amplification in this GTP binding assay, thereby making the derived efficacies more reliable estimates of intrinsic efficacy compared to other assays. Accordingly, the consistently higher efficacies of nitazenes relative to the other opioids underscore their robust capacity to activate MOR despite structural variations at the R1, R2, and R3 positions.

With respect to potency, the GTP binding assay produced a ranking similar to that observed in the cAMP inhibition assay. However, the overall dynamic range of pEC_50_ values was markedly narrower, spanning 1.73 log units compared to 3.33 log units in the cAMP assay. DINZ remained the most potent compound (EC_50_ = 0.39 nM). Unlike in the cAMP assay, the higher-potency nitazenes in the GTP binding assay clustered tightly with DINZ, exhibiting EC_50_ values ranging from 0.52 nM (NP-INZ) to 1.7 nM (INZ). EDZ, IDZ, BNZ, and EP-ENZ formed a lower-potency cluster with fentanyl, near DAMGO (EC_50_ = 8.5 nM), with EC_50_ values spanning from 6.9 nM to 14 nM, at least fourfold higher than those of the higher-potency cluster (Figure 2B).

### Live-cell BRET Go activation confirmed high nitazene efficacy with limited potency separation

While the GTP-G_i_ binding assay yields differential efficacy measurements by minimizing signal amplification, it measures receptor activity in harvested membrane preparations rather than in live cells, thereby limiting extrapolation of these results to MOR function in more native cellular environments. To complement these data and enable comparison under more physiological conditions, we carried out a BRET-based Go activation assay in HEK293T cells transiently expressing MOR (see Methods). Similar to GTP binding, Go activation occurs proximal to receptor activation (Figure 2C), and exhibits lower signal amplification than assays based on secondary messengers.

In this assay, DINZ was greater than threefold more potent than INZ, while DFNZ was approximately twofold more potent than FNZ, and DENZ and ENZ displayed comparable potencies (Figure 2C; Table 1). The R2 N-pyrrolidino compounds (NP-INZ and NP-ENZ) exhibited intermediate potencies between those of their respective N-diethyl and N-desethyl analogues. DINZ remained the most potent compound, with an EC_50_ of 0.54 nM. BNZ, EP-ENZ, and EDZ were less potent than DAMGO (EC_50_ = 1.9 nM); however, the separation between the lower- and higher-potency nitazene clusters was noticeably reduced in this assay compared to both the cAMP inhibition and GTP–G_i_ binding (see Figure 2A-C). Among the fentanyl analogues, CPF (EC_50_ = 1.9 nM) and fentanyl (EC_50_ = 4.2 nM) showed higher potencies, whereas VRF (EC_50_ = 140 nM) was 33-fold less potent than fentanyl.

Similar to observations in the cAMP inhibition and GTP binding assays, all nitazenes displayed efficacies comparable to or exceeding that of DAMGO in this assay, ranging from 99.7% (EDZ) to 127.2% (INZ). Notably, FNZ and DFNZ showed lower efficacies than their isopropoxy and ethoxy counterparts, and all N-desethyl analogues were less efficacious than their N-diethyl counterparts. Morphine and fentanyl also displayed efficacies above that of DAMGO at 106.1% and 116.3%. Buprenorphine and VRF exhibited reduced efficacies of 87.7% and 75.3%, respectively, confirming that this assay adequately differentiated partial from full agonists. Unexpectedly, CPF also demonstrated a reduced efficacy of 89.6%.

### Structural basis of nitazene recognition by MOR and implications for SAR

To interpret the observed SARs in the structural context of MOR, we first examined the recently reported cryo-EM structure of the MOR–G_i_–FNZ complex^24^ and performed a detailed analysis of ligand–receptor interactions. This analysis indicated that FNZ adopts a trivalent binding mode, with the R1, R2, and R3 substituents projecting into spatially distinct receptor subpockets (Figures 3A, 5, and S3). At R2, in coordination with the salt bridge formed between the tertiary amine and the carboxyl side chain of Asp^3.32^, one ethyl substituent on the tertiary amine is accommodated within a subpocket formed by Asp^3.32^, Tyr^3.33^, and Met^3.36^, whereas the other ethyl group is enclosed by Met^3.36^, Trp^6.48^, and Tyr^7.43^. At R3, with the benzimidazole core sandwiched between Ile^7.39^, Gln^2.60^, and Leu^2.57^, the 5-nitro group adopts a parallel orientation relative to the aromatic ring of Tyr^1.39^, consistent with a favorable stacking interaction, and also makes a close polar contact with the side-chain hydroxyl of Tyr^1.39^ (Figure 3A). The 5-nitro group also contacts His^7.36^, as well as the backbones of Ser^2.61^ and Tyr^2.64^. Stemming from the core, the 2-benzyl substituent contacts Trp^7.35^ and Ile^6.51^. At R1, the fluoroethoxy tail extends toward the TM5-TM6 interface and exhibits conformational flexibility, adopting two distinct conformations in the solved structure: one oriented extracellularly towards Lys^5.39^ and Val^5.42^, and the other oriented intracellularly towards Trp^6.48^ and in contact with His^6.52^ (Figure S3).

**Figure 3:**
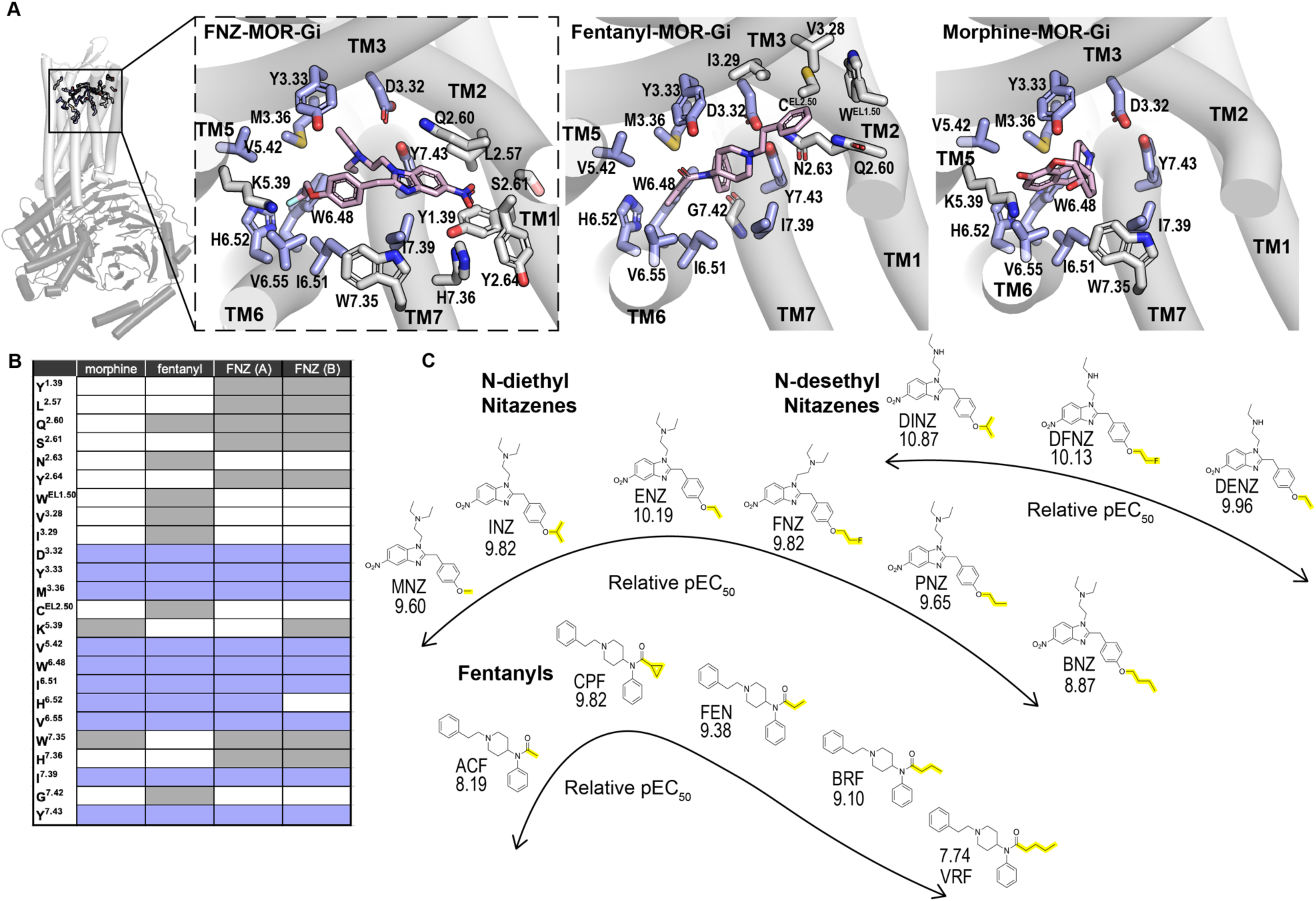
Comparison of binding poses and functional potency across opioid classes. (A) Cryo-EM structures of ligand-bound MOR-G_i_ complexes for FNZ (PDB: 9O36),^24^ fentanyl (PDB: 8EF5), and morphine (PDB: 8EF6).^25^ MOR residues within 5 Å of each indicated ligand are shown in (A) and listed in (B). Residues highlighted in periwinkle represent contacts shared by all three ligands, defining the MOR orthosteric binding site (OBS).^34^ (C) Structural and functional comparison of N-desethyl nitazenes, N-diethyl nitazenes, and fentanyl analogues with different R1 groups (defined in Figure 1). The listed potencies are the mean pEC_50_ values for the indicated compounds from cAMP inhibition experiments in this study (Table 1) and our previous work.^13^ Yellow highlights differences in chain length among the compounds, as well as the region of each molecule that occupies the interface of TMs 5, 6, and 7 within the OBS. Notably, for both scaffolds, functional potency is optimized in compounds with two carbons extending beyond the oxygen. However, removal of an ethyl group from R2 alters the relationship between chain length and potency, as demonstrated in the diverging trend for N-desethyl nitazenes compared to N-diethyl nitazenes and fentanyls. Specifically, within the N-desethyl series, DINZ is the most potent analogue, and DFNZ is more potent than DENZ despite having one additional heavy atom in its alkoxy chain.

In comparison with the fentanyl (PDB: 8EF5) and morphine (PDB: 8EF6) binding poses with MOR,^25^ we found, as expected, that these three opioid scaffolds commonly occupy the orthosteric binding site (OBS) enclosed by TMs 3, 5, 6, and 7,^34^ and share most of the R1 and R2 contact residues (Figure 3A–B). However, except for interaction with Trp^7.35^ and Lys^5.39^, morphine does not extend much beyond the OBS, while the phenylethyl moiety of fentanyl protrudes into the interface between TMs 2 and 3 but does not reach the R3 subpocket at the interface among TMs 1, 2, and 7, as does the 5-nitro group of FNZ (Figure 3).

Our binding pocket analysis further revealed an intramolecular hydrophobic-aromatic interaction between the two R2 ethyl groups (attached to the tertiary amine) and the 2-benzyl substituent (Figures 3A and 4A), which appears to pre-organize these moieties for optimal accommodation within the OBS. In contrast, no comparable intramolecular interaction was observed in the MOR-bound poses of fentanyl and morphine. However, this intramolecular interaction in FNZ was not readily apparent and could reflect either an intrinsic conformational preference of the free ligand, receptor-induced stabilization, or a feature unique to FNZ relative to other nitazenes.

**Figure 4:**
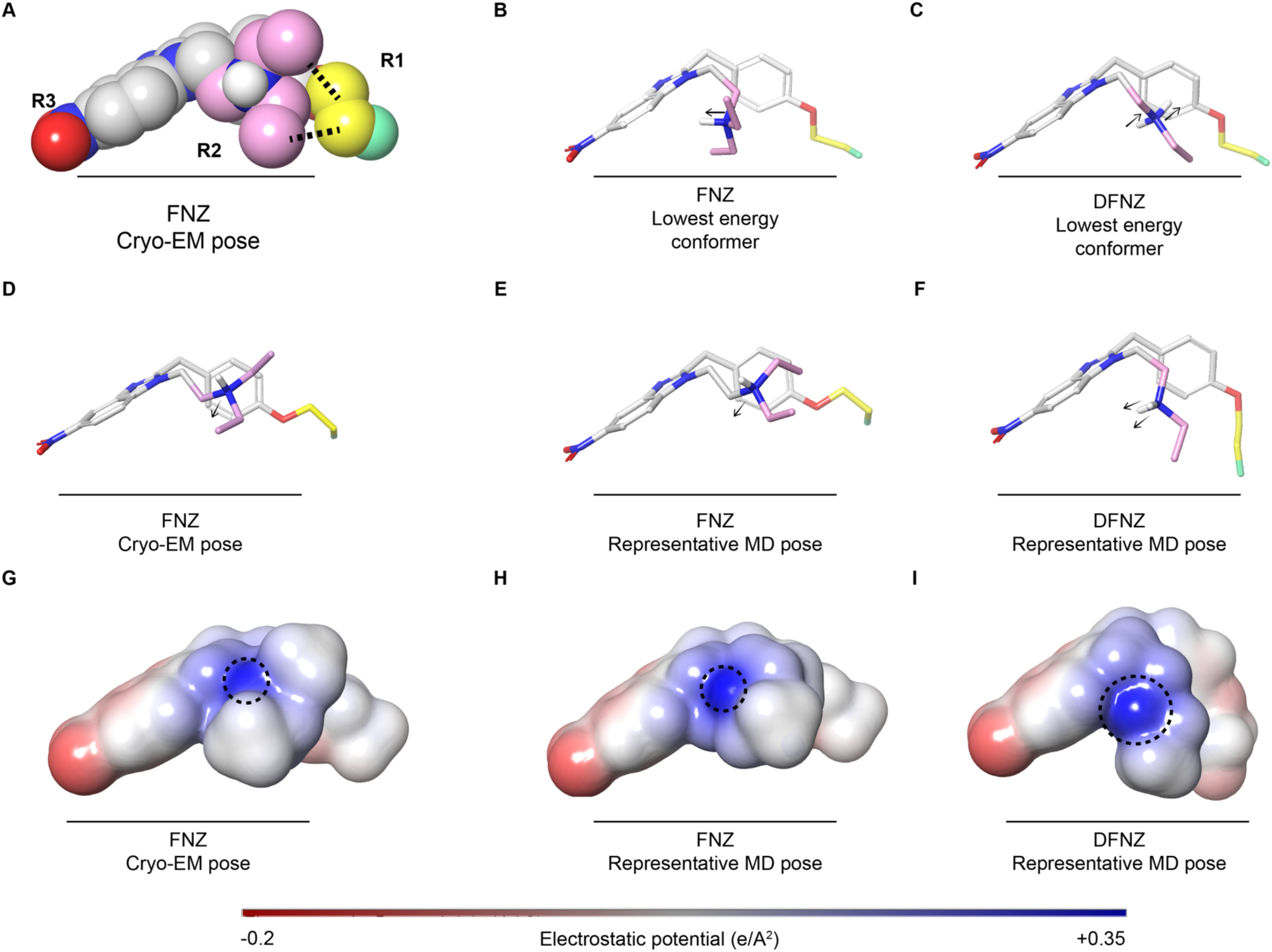
Conformational and electrostatic variation between nitazene analogues and lowest-energy vs. MD-simulated representative conformers. (A) FNZ in spherical representation, shown in its cryo-EM pose (PDB: 9O36).^24^ Dashed lines indicate intramolecular interactions. (B,C) Lowest-energy conformers of FNZ (B) and DFNZ (C), identified by molecular mechanics-based conformational searches. (D) The cryo-EM pose of FNZ shown in (A), displayed in stick representation and from the same viewing angle. (E,F) Representative MOR-bound poses of FNZ (E) and DFNZ (F) from MD simulations, viewed from the same angle as in (D). Arrows in (B-F) indicate the orientation of the proton on the charged R2 amine, which differs systematically between the lowest-energy conformers of the free ligands (B,C) and the corresponding MOR-bound poses (D-F). Yellow and pink denote atoms belonging to the R1 and R2 moieties, respectively, as defined in Figure 1. (G-I) QM-calculated electrostatic potential surfaces for FNZ in its cryo-EM pose (G) and its MD pose (H), and for DFNZ in its MD pose (I). Dotted circles mark the distribution of positive surface charge, which is widest in DFNZ.

**Figure 5:**
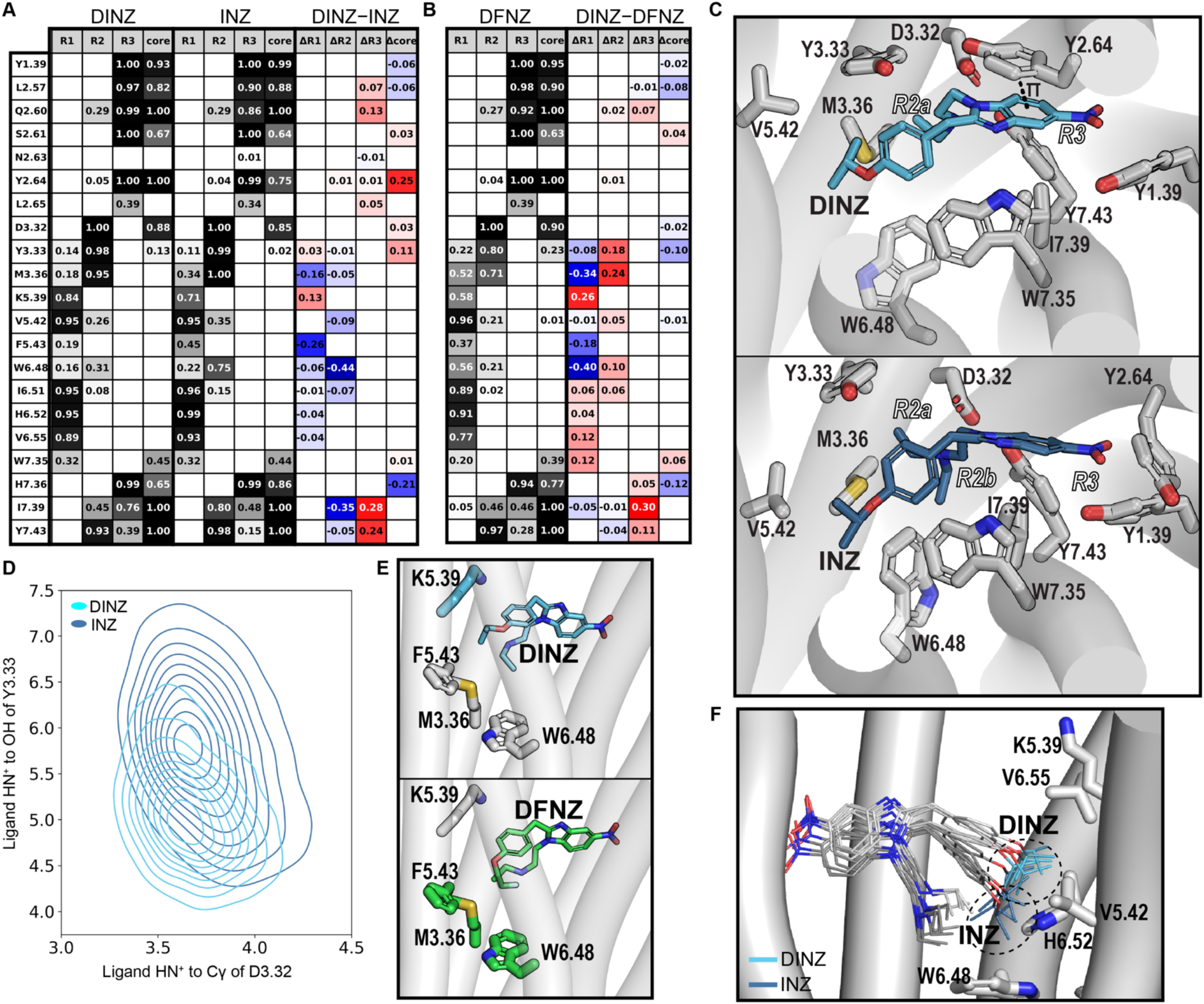
DINZ uniquely coordinates R1, R2, and R3 subpockets. (A,B) Comparison of regional contact frequencies for DINZ versus INZ (A) and DINZ versus DFNZ (B). Ligand region definitions are shown in Figure 1. Contact is defined as any residue heavy atom within 5 Å of the indicated ligand region. Individual contact frequencies for each ligand are shown in grayscale, and differences in contact frequency are shown on a red-to-blue scale, where red denotes positive values and blue denotes negative values. (C) Comparison of the R2 and R3 binding poses of DINZ (top) and INZ (bottom). R2: the ethyl group of DINZ remains confined to the R2a subpocket, formed by TM3 residues and Tyr328^7.43^, whereas in INZ, one ethyl occupies the R2a subpocket and the other extends into a second subpocket formed by TM6 and TM7 residues (R2b). R3/core: DINZ and other N-desethyl compounds show higher contact frequency with Tyr130^2.64^ (pink) than N-diethyl compounds, promoting a π-stacking interaction between the ligand core and the tyrosine side chain. In contrast, N-diethyl compounds show reduced engagement with Tyr130^2.64^, resulting in the rotamer shift observed for this residue in INZ (bottom). (D) The distribution of the distances from the protonated R2 amines of DINZ and INZ to Cγ of Asp149^3.32^ and OH of Tyr150^3.33^. Although both compounds show similar average distances to Asp149^3.32^, DINZ binds approximately 1 Å closer to Tyr150^3.33^ and exhibits a narrower distance distribution to both residues than INZ. (E) Comparison of R1 interactions in DINZ (top; cyan) and DFNZ (bottom; green). The R1 tail of DINZ is oriented toward the extracellular side. Residues shown in cyan indicate higher contact with the R1 tail of DINZ, whereas residues shown in green indicate higher contact with the R1 tail of DFNZ. (F) Comparison of the R1 tail distribution of DINZ (white and cyan) and INZ (gray and dark blue) in equilibrated trajectories. Although these compounds share an R1 isopropoxy tail, their R1 moieties occupy distinct spatial distributions within the binding pocket. DINZ is oriented extracellularly toward Lys235^5.39^, whereas INZ is oriented intracellularly toward His299^6.52^.

While the cryo-EM structure provides a high-resolution snapshot of FNZ bound to MOR, a static structure cannot fully delineate the receptor structure-based SARs observed across the nitazene scaffold, particularly given the conformational flexibility of the R1 and R2 substituents and their potential allosteric coupling within the receptor. In the following sections, we describe the distinct structural and dynamic features of these analogues, both as free ligands and when bound within the MOR binding pocket.

### Free and MOR-bound nitazenes exhibited similar conformations but differed in R2 orientation

To distinguish between conformational preferences of the free ligand and receptor-induced stabilization within MOR, we performed an exhaustive conformational search to characterize the conformational landscape of selected nitazene analogues in their unbound state (see Methods). The conformational search employs molecular mechanics-based energy minimization to systematically sample conformational space and identify low-energy conformers for each ligand.^35^ Given the multiple rotatable bonds within the nitazene scaffold, these compounds are highly flexible and sample a wide range of conformations in the search. Notably, intramolecular interactions are a shared feature among nitazenes and stabilize a subset of these conformations in aqueous solution, including the lowest-energy conformers.

By comparing the lowest-energy FNZ conformer yielded by the search with the MOR-bound FNZ pose revealed by the cryo-EM structure, we found that while both have similar intramolecular interactions between the R2 ethyl groups and the 2-benzyl ring, the orientation of the R2 N-H bond differs significantly between the lowest-energy conformer of free FNZ and the experimentally determined MOR-bound FNZ pose (Figures 4B-4F and S4). In the MOR binding site, the tertiary amine forms a critical salt bridge with residue Asp149^3.32^, constraining the N-H bond to orient away from the ligand core (Figure 4D-4F). In contrast, the R2 N-H bond in the lowest-energy conformer of FNZ is oriented inward towards the ligand core (Figure 4B-4C and S4). Thus, formation of the salt-bridge interaction with Asp149^3.32^ must offset the energetic penalty associated with reorienting the N-H bond in the MOR-bound conformation.

### R2 N-desethylation and R1 tail variation altered intramolecular interactions and electrostatic properties

Among the lowest-energy conformers of the free nitazene analogues, the orientation of the R2 amine and its substituents also differed markedly between the corresponding N-diethyl and N-desethyl analogues. In the N-desethyl analogues, the protonated amine bears two N-H bonds, both oriented inward towards the alkoxybenzyl substituent, consistent with an intramolecular cation-π interaction in which the positively charged nitrogen engages the aromatic π system of the 2-benzyl substituent, thereby stabilizing the ligand conformation (Figure 4C). In contrast, in the N-diethyl analogues, the bulky ethyl substituents reorient the R2 amine such that the remaining N-H bond points towards the benzimidazole core, potentially interacting with its π-orbitals. The two ethyl chains fan out around the nitrogen, positioning one ethyl to interact with the 2-benzyl ring, consistent with a CH-π interaction in which the hydrogens of the ethyl group weakly interact with the aromatic π face of the ring (Figure 4B). The second ethyl strays outwards, away from the ligand core.

In all N-desethyl compounds, INZ, and BNZ, one R2 ethyl group lies within 4.9 Å of the R1 terminal heavy atom (shortest in BNZ at 3.9 Å), whereas FNZ and ENZ do not exhibit this proximity (6.9 Å and 7.2 Å, respectively). Furthermore, the branched isopropoxy (INZ) and elongated butoxy (BNZ) R1 tails position the R1 terminus closer to R2, compared to ethoxy (ENZ) or fluoroethoxy (FNZ) tails, demonstrating that increasing the hydrophobic surface area of the alkoxy chain, in addition to N-desethylation, also increases R1-R2 intramolecular interaction in free nitazenes. In BNZ, the short R1-R2 distance is associated with an inward-facing N-H bond orientation that engages the R1 moiety, resembling that observed in the N-desethyl compounds (Figure S4).

To assess how conformational differences influence the electrostatic properties of each analogue, we performed quantum-mechanical (QM) calculations to characterize electrostatic potential (ESP) surfaces (see Methods) for FNZ and DFNZ in both their lowest-energy conformers and their MOR-bound poses. The MOR-bound poses displayed broader distributions of the positive charge on the R2 surface than the lowest energy conformers extracted from the conformational search. Notably, DFNZ exhibited a markedly larger positive surface area at R2 upon MOR engagement compared to FNZ, consistent with greater accessibility of the N-desethyl nitrogen for interaction with Asp149^3.32^ in the MOR binding site (Figure 4G-4I). This finding suggests that N-desethylation results in a more favorable ESP surface for recognition by Asp149^3.32^, which is consistent with our experimental results, where DINZ and DFNZ were more potent than their N-diethyl analogues across all three assays. However, as DENZ remained less potent than ENZ in both the cAMP inhibition and GTP-G_i_ binding assays, this change in ESP contributes only partially to the nitazene SAR.

### MD simulations revealed conserved trivalent binding architecture with analogue-specific R1-R2 coupling

To evaluate how these differences in conformations and intramolecular interactions influence the binding of nitazene analogues at MOR, we performed extensive simulations of MOR in complex with selected nitazenes tested in this study, in an explicit lipid bilayer–water environment, based on the MOR–G_i_–FNZ cryo-EM structure (see Methods; Table S1).

For the MOR–G_i_–FNZ complex, we initiated parallel simulations with FNZ in both R1 tail poses revealed by the cryo-EM structure (poses A and B; see Figure S3), each with multiple replicas. The overall receptor remained stable in the lipid bilayer–water environment and largely preserved the cryo-EM conformation across all trajectories. During equilibration, the FNZ R1 tail converged to a conformation closer to the pose A position, with the terminal -F in contact with His299^6.52^ and Val302^6.55^, and oriented downwards towards Trp295^6.48^ (Figure S3). Notably, the tail demonstrated local flexibility within this subpocket, sampling multiple orientations.

Across the simulations of all complexes, the benzimidazole core maintained a consistent orientation within the binding site, with the planar faces of its π system interacting with Ile324^7.39^ and Gln126^2.60^, as observed in the FNZ bound MOR cryo-EM structure. However, modifications at R1 and R2 altered the orientations adopted by R1, R2, and R3 within their respective subpockets. To characterize the molecular determinants underlying these substituent-dependent effects, we identified ligand-contacting residues for all nitazene analogues and quantified their contact frequencies in the MD simulations for each complex (see Methods). This analysis identified 21 residues that interacted with at least one analogue (Table 2). Notably, 17 of the 18 residues within 5 Å of FNZ in the cryo-EM structure (listed in Figure 3B) exhibited contact frequencies greater than 0.30 with FNZ in the simulations, supporting the stability of the binding pose and the reliability of our simulation system (Table 2). To further dissect ligand-residue interactions, we partitioned each ligand into R1, R2, R3, and benzimidazole core regions (see Figure 1A for region definition) and calculated region-specific contact frequencies. This analysis showed that all analogues maintained the trivalent binding mode throughout the simulations, though different analogues exhibited distinct contact patterns within each subpocket (Figures 5A-5B and S5).

An examination of the R2 region showed that in the N-diethyl analogues, the two R2 ethyl groups occupy two adjacent subpockets, with each ethyl residing in a distinct subpocket (Figure 5C). In contrast, in the N-desethyl counterparts, the single remaining R2 ethyl shifts toward an altered position, resulting in reduced contact frequencies with Trp295^6.48^, Ile298^6.51^, Ile324^7.39^, and Tyr328^7.43^, and forming stronger contact with Tyr150^3.33^ (Figures 5A, 5C, S5A, S5D). Specifically, the amine of N-desethyl analogues bound nearly 1 Å closer to Tyr150^3.33^ than N-diethyl analogues, suggesting that N-desethylation allows the remaining ethyl group to form a tighter interaction with Tyr150^3.33^, although the salt-bridge distance to Asp149^3.32^ did not differ between the two series (Figure 5D). Notably, among all nitazene analogues, DINZ exhibited the most compact R2–TM3 interaction (Figure 5D), consistent with its highest potency. Compared to DINZ, the amine interactions of DENZ at Asp149^3.32^ and of DFNZ at Tyr150^3.33^ were more dynamic (Figures 5D,S5B, and S5E), and a similar sensitivity to R2 orientation was observed for EP-ENZ, where stereochemistry-dependent positioning of the R2 amine weakened its interaction with Asp149^3.32^ (see Supporting Results and Figure S6).

These R2 changes propagated to enable the benzimidazole core of the N-desethyl analogues to interact more frequently with Tyr130^2.64^, consistent with a stronger propensity for π-stacking with this residue (Figure 5C), while their R3 5-nitro group attached to the core engaged residues Ile324^7.39^ and Tyr328^7.43^ with higher contact probability. These differences are consistent with reduced steric repulsion near TMs 2 and 7 upon removal of an R2 ethyl group, which allows closer engagement between these residues, the ligand core, and R3 due to the repositioning of the remaining R2 ethyl group. Notably, a similar correlation is observed in the comparison between the two N-desethyl analogues DINZ and DFNZ, where DINZ, the more potent analogue, exhibits stronger engagement at these residues. This observation suggests that variation at R1 can likewise propagate to influence R3 interactions (Figure 5).

As the absence of the R3 5-nitro group markedly reduced the experimental potency, as demonstrated by the comparison of IDZ with INZ and EDZ with ENZ, it is tempting to speculate that the coordinated changes propagated from R2 N-desethylation to R3 may contribute to the higher potency of DINZ compared to INZ, and to a lesser extent, DFNZ than FNZ (Figure 2 and Table 1). However, in the ENZ and DENZ pair, although they exhibit divergent R3 engagements similar to the other pairs, the N-desethyl analogue DENZ has comparable to, or even slightly lower potency than, ENZ (Table 1). The differences among these pairs are therefore likely influenced by both R3 engagement and distinct configurations within the R1 subpocket (see below).

In addition to enhancing R3 engagement with the receptor across the N-desethyl analogues, the repositioning of the R2 ethyl enabled increased R1 contact with Lys235^5.39^ in DFNZ and DINZ compared to their N-diethyl counterparts, FNZ and INZ, respectively. As shown in Figures 5F and S5F, this increase arises because N-desethylation shifts the spatial distribution of the R1 tail within the subpocket in the FNZ/DFNZ and INZ/DINZ pairs. Within the N-desethyl series, DINZ and DFNZ exhibit distinct R1 spatial distributions (Figure 5E, 5F, and S5F), reflecting differences in R1 substitution. This divergence in R1 positioning further propagates to the R3 subpocket, as discussed above. In contrast, ENZ and DENZ did not display a comparable difference in R1 contact frequencies, despite their analogous relationship to the FNZ/DFNZ and INZ/DINZ pairs (Figure S5). Indeed, for the ENZ/DENZ pair, the R1 tails adopt the same spatial distribution despite the difference in R2 and R3 engagements (Figure S5C).

Together, these findings indicate that R1, R2, and R3 exhibit coordinated, context-dependent coupling within the MOR binding pocket. N-desethylation at R2 alters the positioning of the remaining ethyl group, which in turn reshapes R1 spatial occupancy and R3 engagement, while variation at R1 also modulates the R3 engagement. The conformational configurations resulting from this three-way allosteric interplay differ among the fluoroethoxy, isopropoxy, and ethoxy analogue pairs, and likely underlie the divergent impact of N-desethylation observed experimentally, where the potency shift was greatest in the isopropoxy series.

### Varied R1 engagement associates with differences in both potency and efficacy

Our experimental results showed that extending the R1 ethoxy chain of ENZ to the butoxy chain of BNZ resulted in a 20-fold decrease in potency in the cAMP inhibition assay, along with 8.7% and 21.1% reductions in efficacy in the GTP binding and Go activation assays, respectively (Table 1). Because the R1 chain directly interacts with TM6, whose conformational changes determine receptor (de)activation, and by extension, ligand efficacy, we sought to investigate how variations in the R1 alkoxy substituent affect the conformation at the TM5 and TM6 interface to connect efficacy with ligand-receptor interactions.

Specifically, we measured the distance between the Cα atoms of Val238^5.42^ and Val302^6.55^ in the extracellular portions of the TM5 and TM6, respectively, during the simulations.^30^ Among all simulated nitazenes, DINZ maintained the most stable and narrowly distributed Val238^5.42^–Val302^6.55^ distance, centered near 12 Å (Figure 6A), suggesting optimal complementarity with the binding pocket that may contribute to its highest potency. In contrast, compounds with lower potency exhibited broader and multimodal distance distributions, indicating reduced pocket complementarity and increased binding pocket flexibility. In the BNZ-bound complex, the BNZ butoxy tail protruded deeper into the cavity between TM5 and TM6, inducing an outward shift in TM6 (Figure 6B).

**Figure 6:**
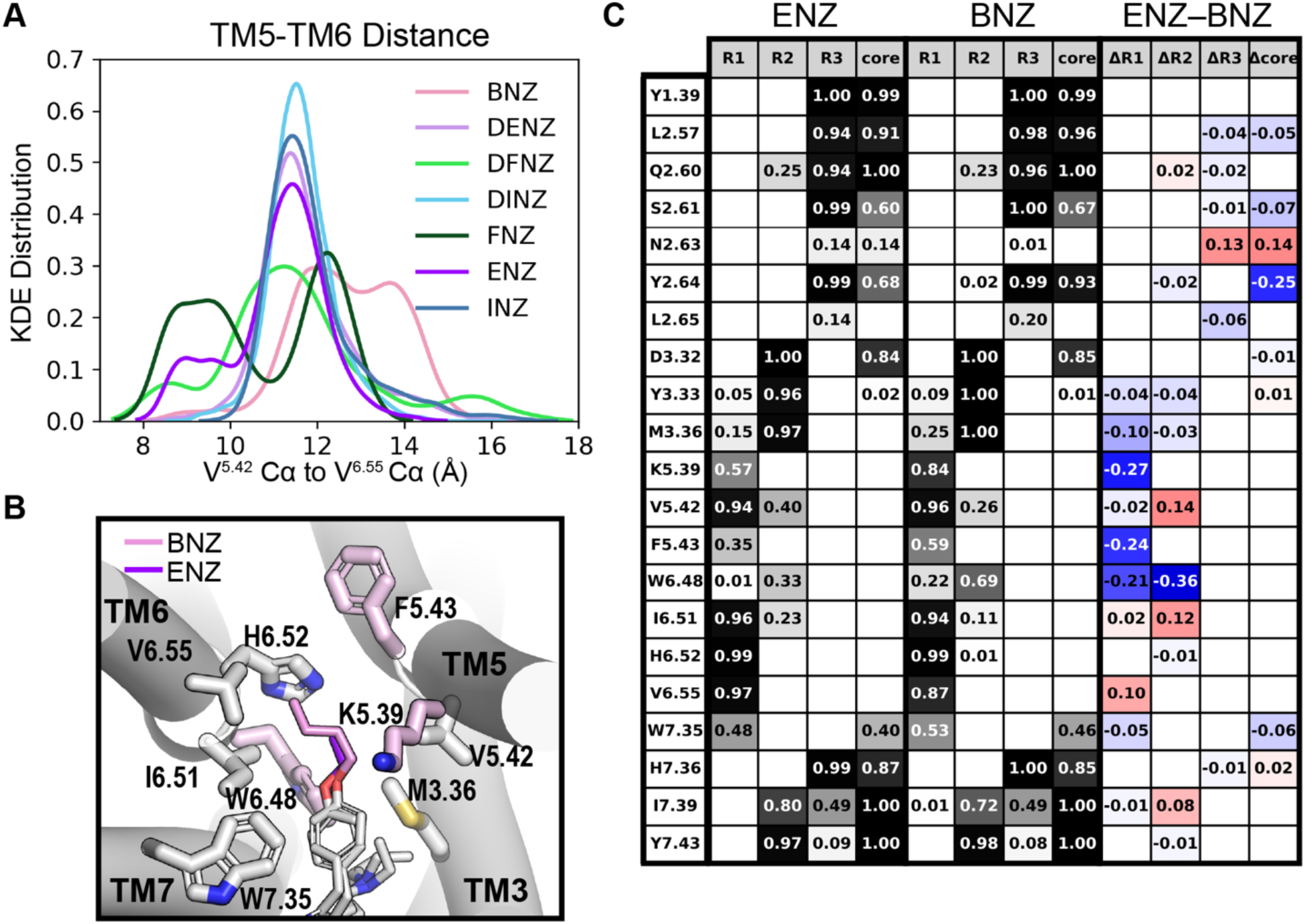
R1 variation shifts conformation of TM6. (A) The distribution of the Cα-Cα distance between Val238^5.42^ and Val302^6.55^ in MD simulations. (B) The differences in R1 contacts between ENZ and BNZ. Light pink residues indicate >10% higher contact frequency with the R1 group of BNZ (pink) than with that of ENZ (purple). (C) Comparison of regional contact frequencies between ENZ and BNZ using the same protocol as in Figure 5 and described in the Methods.

To further probe how R1 substituent variation influences the TM5-TM6 interface, we compared the contact frequencies of ENZ and BNZ (Figure 6C). Despite BNZ being substantially less potent than ENZ, its contact frequency with Lys235^5.39^ was higher than ENZ and reached 0.84, matching that of DINZ. However, visual inspection revealed that BNZ’s interaction with the R1 subpocket differs substantially from DINZ and ENZ: while the terminus of DINZ remained localized near Lys235^5.39^ and that of ENZ is limited to His299^6.52^, resulting in consistent and stable interactions, the butoxy tail of BNZ, owing to its bulk and flexibility, perturbs the TM5–TM6 interface. Specifically, proximal carbons of the BNZ tail engage TM5 near Lys235^5.39^, while distal carbons extend downwards towards TM6, increasing contact with Trp295^6.48^ and Phe239^5.43^ compared to ENZ (Figure 6B). This broader distribution of contacts suggests that BNZ does not form a stable, localized interaction but instead disrupts the local packing within the subpocket.

Although this disruption at the R1 subpocket likely propagates to affect TM6 movement, resulting in reduced efficacy, BNZ remains highly efficacious and potent at MOR, exhibiting higher efficacy and potency than fentanyl in the GTP binding assay. We propose that this preserved activity arises from tight engagement of the main scaffold with MOR. In particular, compared to ENZ, BNZ showed increased contact with Tyr130^2.64^, and maintained the tight π-stacking interaction between the benzimidazole core and phenol sidechain observed for DINZ (Figure 5C).

Together, these results show that R1 substituents modulate potency and efficacy by altering conformational behavior at the TM5–TM6 interface. Optimal R1 engagement, as in DINZ, stabilizes the pocket and supports efficient receptor activation, whereas bulkier groups such as the butoxy chain in BNZ disrupt local packing.

## DISCUSSION

The differences observed across our *in vitro* assays reflect their complementary readouts, with the cAMP inhibition assay providing high sensitivity for potency ranking, the GTP–G_i_ binding assay offering a lower-amplification measure of intrinsic efficacy, and the BRET Go activation assay capturing receptor activity in a live-cell context. Across all assays, nitazenes consistently exhibited potency comparable to or greater than the full agonist reference, DAMGO. This finding aligns with previous reports^5, 12, 13, 18, 22, 36^ and underscores the profound public health risk these substances pose.^10, 16, 37^ The trend was particularly evident in the cAMP inhibition assay, a robust measure of ligand potency,^13^ where all tested nitazenes, along with fentanyl and its analogue CPF, displayed higher pEC_50_ values than DAMGO. While fentanyl and CPF grouped with the nitazenes in this assay, their potencies were lower than the more potent 5-nitro nitazenes.

Our *in vitro* work elucidated the SARs governing the high potency of nitazenes by identifying three substituent modifications that reduced potency consistently across the assays: steric bulk at R1 (BNZ) or R2 (EP-ENZ) position and the absence of a 5-nitro group on the benzimidazole core (IDZ and EDZ). While previous studies have shown that direct halogenation of the R1 position severely diminishes potency,^12, 14, 38^ our investigation of the fluoroethoxy substituted analogues FNZ and DFNZ showed that both analogues retained high potency, likely due to preservation of the alkoxy chain and optimal R1 chain length. The reduced adverse effects of FNZ and DFNZ *in vivo* have spurred investigation into their therapeutic potential as strong analgesics with an improved safety profile.^24^

While modifications at all three pharmacophoric positions influenced potency, R1 modifications exerted the most pronounced effect on efficacy. Specifically, INZ consistently showed the highest experimental efficacy, ranking as the single most efficacious compound in two of the three assays. Conversely, the butoxy R1 tail in BNZ reduced efficacy, most prominently in the GTP-G_i_ binding assay where BNZ was among the few nitazenes with efficacy below that of DAMGO, consistent with steric constraints imposed by the larger substituent. Finally, although fluoroethoxy R1 chains preserved potency, FNZ and DFNZ exhibited slightly reduced efficacies, particularly in the BRET Go activation assay, where both fluorinated analogues demonstrated lower efficacies than unfluorinated analogues with the same R2 substituent, ENZ and DENZ, respectively.

Structure-based computational approaches are instrumental for characterizing ligand-receptor recognition. However, until the recent resolution of the MOR–Gi–FNZ structure, such studies of nitazene recognition at MOR were limited by the lack of an experimentally determined nitazene-bound MOR structure. Notably, however, de Luca et al. complemented their experimental study of nitazenes with molecular modeling and simulations, finding that INZ and metonitazene formed more stable complexes with MOR than fentanyl.^39^ Clayton et al. employed the MOR structures complexed with several non-nitazene opioids as templates in their modeling work, and identified an ENZ pose consistent with the recently resolved MOR structure bound with FNZ.^40^ Nevertheless, no prior modeling study has systematically characterized MOR structure-based SAR across scaffolds.

Our integrated conformational, electrostatic, and MD analyses indicate that the nitazene SAR at MOR arises not simply from additive substituent effects, but also from coordinated intraligand and ligand–receptor coupling across the R1, R2, and R3 regions. Although all analogues retained a conserved trivalent binding architecture, MD simulations revealed substituent-dependent differences in subpocket engagement. Our MD results showed that the persistent but differential occupancy of the R1 subpocket by nitazenes, where they interact with TMs 5 and 6, may relate to the differences in efficacy in addition to potency. In particular, the bulky butoxy tail of BNZ induced a greater TM5-TM6 distance, potentially indicating a trend towards an inactive state (Figure 6A-B), as it may hinder the conformational changes of TM6 during MOR activation required to create the binding interface for G protein coupling.^41^ Notably, this finding is consistent with our previous work comparing valerylfentanyl and fentanyl, in which the bulky N-pentanamide tail of valerylfentanyl caused a similar outward motion of TM6.^30^ Thus, our simulation results explain the optimal R1 chain lengths and shapes observed *in vitro* for both the fentanyl and nitazene scaffolds—namely, isopropoxy and ethoxy for nitazenes, and cyclopropyl for fentanyls—and highlight a relatively unified SAR at the R1 subpocket (Figure 3C).

However, for nitazenes, the R2 substituent plays a substantial role in shaping receptor engagement alongside R1. Our MD results revealed that N-desethylation altered the intramolecular interaction network and increased the electrostatic accessibility of the protonated R2 amine. QM analysis further showed broader positive electrostatic surfaces upon MOR binding in the N-desethyl analogues, consistent with tighter interactions near Asp149^3.32^ and Tyr150^3.33^. The resulting positional shift propagated to reshape both R1 engagement and R3 interaction in a substituent-dependent manner, supporting coordinated coupling among R1, R2, and R3 within the MOR binding pocket. Specifically, in the fluoroethoxy and isopropoxy pairs, N-desethylation at R2 allosterically strengthened both R1–receptor interactions and R3 engagement, whereas this coordination is absent in the ethoxy pair, providing a structural basis for the 11-fold increase in potency of DINZ relative to INZ, compared with the similar potency between DENZ and ENZ in the cAMP inhibition assay. Thus, the exceptional potency of DINZ arises from coordinated optimization of R2–TM3 engagement, TM5–TM6 stabilization, and R3 engagement, yielding a “sweet spot” of near-perfect geometric and electrostatic complementarity between ligand and receptor within the MOR binding site.

Based on our combined experimental and simulation findings, we propose a nitazene SAR at MOR that differs from the fentanyl SAR by engaging the R3 subpocket and by allosterically coupling R1 and R2 interactions. This distinct SAR provides a structural framework for understanding the unusually high potency and efficacy of nitazenes and may aid in predicting the activity of newly emerging analogues in illicit drug markets.

## METHODS

### HTRF-Based cAMP assay

Cell culture condition and assay protocol were adopted from Tsai et al.^13^ Briefly, the inhibition of forskolin-stimulated cAMP accumulation by MOR agonists was assessed using the Revvity Homogenous Time-Resolved Fluorescence (HTRF) cAMP G_i_ kit (Revvity, Waltham, MA, USA). This kit uses a Förster resonance energy transfer (FRET) signal between two fluorescent dyes to measure the cAMP levels. Specifically, the cAMP produced by cells competes with europium-cryptate-labelled cAMP (donor) for binding to the d2-labeled anti-cAMP antibody (acceptor). The concentration of unlabeled cAMP produced by the Gi/o pathway of cells is inversely proportional to the FRET signal: the higher the FRET signal, the lower the concentration of unlabeled cAMP produced.

The FLP-FRT-HEK cells stably expressing the human MOR were grown to 80% confluency in Dulbecco’s Modified Eagle Medium (DMEM) containing 10% fetal bovine serum, 2 mM L-glutamine, 1% penicillin-streptomycin, and 50 μg/mL hygromycin B. On the day of the experiment, cells were washed three times with phosphate-buffered saline (PBS) buffer and dissociated from cell culture dishes using 0.05% trypsin-ethylenediaminetetraacetic acid (EDTA) before being centrifuged at 1000 rpm for 5 min. The collected cell pellet was suspended in stimulation buffer 1 from the kit, and 3000 cells/well (5 µL) were transferred to HTRF 96-well low volume plates (Revvity). The cells were then incubated with 4 µL of test compounds at indicated concentrations for 10 minutes at 37°C. Afterward, 1 µL of forskolin (50 µM) was added, and the cells were incubated at 37°C for 45 min. Finally, 5 µL of europium-cryptate-labelled cAMP (Eu-cAMP) and 5 µL d2-labelled antibody were added, and the cells were incubated at room temperature for 1 hour.

The FRET signal level was assessed by calculating the fluorescence ratio of 665 and 620 nm emissions, which were measured using the PHERAstar FSX plate reader (BMG Labtech, Cary, NC, USA), with the “integration start” and the “integration time” of the HTRF optic module set to 60 and 400 µs, respectively.

All compounds were tested at the specified concentrations in at least four independent experiments (n ≥ 4), with triplicate wells for each concentration. The E_max_ and pEC_50_ shown in Table 1 were derived by fitting the data to “log(agonist) vs response (three parameters)” dose-response model implemented in GraphPad Prism (version 10.5).

### HTRF-based GTP-G_i_ assay

This assay used the ValiScreen Human Opioid Mu (OP3) CHO-K1 host cell line, which stably expresses the human MOR (Revvity, Waltham, MA, USA). Cells were grown in Gibco Ham’s F-10 Nutrient Mix, supplemented with 10% Fetal Clone II, 1 mM sodium pyruvate, and 250 µg/ml hygromycin B.

To harvest the membrane proteins from the stable cell line, the cells were first grown for 48 hours in 10 cm cell plates until reaching approximately 80% confluence. Cells were detached using Earle’s Balanced Salt Solution (EBSS) containing 5 mM EDTA, and transferred to 50 mL tubes for centrifugation at 4000 rpm for 10 minutes at 4°C. The resulting pellets were resuspended in lysis buffer (50 mMTris HCl, 320 mM sucrose, 2 mM EDTA, 5 mM MgCl_2_). The mixture was vortexed until cells were completely lysed. Subsequently, the lysed cells underwent centrifugation at 10,000 rpm for 30 minutes at 4°C. The resulting pellet was resuspended in the stimulation buffer (Buffer 3) from Revvity’s HTRF GTP/G_i_ kit. The suspension underwent five rounds of homogenization using a Polytron PT 2500 E Homogenizer at 10,000 rpm for 15 seconds, with the samples kept on ice between cycles to prevent overheating. Protein concentration was determined using Pierce BCA Protein Assay Kit (ThermoFisher Scientific Inc.) following the manufacturer’s instructions. The protein preparation was then diluted to the desired concentration with Revvity’s stimulation buffer and stored in aliquots at -80°C.

To measure G_i_ protein activation, the HTRF GTP/G_i_ assay (Revvity, Waltham, MA, USA) employs europium-cryptate-labelled GTP that replaces the unlabeled GDP bound to the Gα_i_ subunit upon receptor activation. When europium-cryptate-labelled GTP (Eu-GTP) binds to Gα_i_, a Förster resonance energy transfer (FRET) signal is produced as energy is transferred from Eu-GTP (donor) to a nearby d2-labelled anti-G_i_ antibody (acceptor) bound nearby. The strength of the emitted FRET signal is proportional to the extent of agonist-induced receptor activation.

The stimulation buffer used in this study was supplemented with MgCl_2_ to a final optimized concentration of 50 mM. Test compounds were diluted in this buffer to 4 times their final concentrations, then dispensed as 5 µL aliquots into HTRF 96-well low volume plates (Cisbio, PerkinElmer). Detection reagents, Eu-GTP and d2-labelled anti-G_i_ antibody, were also diluted to 4 times their final concentrations and added as 5 µL aliquots to each well. 2.5 µg of protein was then added in 5 µL of buffer to each well, yielding a final assay volume of 20 µL per well. The plates were sealed and incubated at 37°C for 24 hours.

The FRET signal level was quantified by calculating the fluorescence ratio of emissions at 665 and 620 nm, measured on a PHERAstar FSX plate reader using the HTRF optic module, with the “integration start” and “integration time” set to 60 and 400 µs, respectively.

While GTPγS and GTP-G_i_ assays are valuable for distinguishing full from partial agonism, conducting these assays at MOR poses a technical challenge. The phenomenon of the “bell-shaped” dose-response curve is well documented^33^ but is often not addressed.

This bell-shaped pattern complicates accurate EC_50_ determination using conventional sigmoidal models, as decreasing responses at higher concentrations obscure the true maximum efficacy of the system. Although GDP and NaCl titration can reduce the bell-shaped characteristic, these reagents effectively “cap” the efficacy of the system by promoting a shift in MOR conformation from its active to inactive form.^42^

To address this issue and obtain more reliable quantitative estimate of efficacy, we first fit the GTP-G_i_ data with a bell-shaped model and, when that model was not appropriate, with a conventional sigmoidal model. Specifically, we first employed the bell-shaped dose-response curve fitting model available in GraphPad Prism, which is defined by the following equations (see Figure S2 for a visual representation of the model parameters):

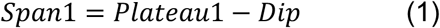

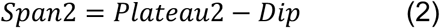

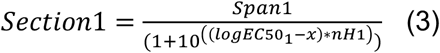

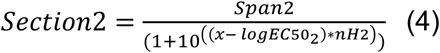

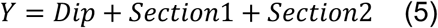

However, we found that the default initial guesses and constraints in Prism were not well suited to our dataset, because fitting the descending curve on the right (dotted gray in Figure S2B) required substantially more datapoints at higher drug concentrations, which could not be tested. Although more extensive parameterization within Prism could substantially improve the fit, doing so required considerable manual effort. To streamline this process through an automated procedure, we implemented the model in Python with the lmfit package, allowing the parameters to be optimized more efficiently for our dataset, with the following consideration.

In our current study, accurate fitting of the ascending curve on the left side (defined by plateau_2, logEC50_2, and dip) was more important than that of the descending curve on the right side (plateau_1, logEC50_1, and dip), because our goal was to capture the primary ligand effect rather than potential off-target effects at higher concentrations. Based on this rationale, we defined a set of constraints to preserve the integrity of ascending curve when the two curves competed to account for the datapoints (see the parameters in the Jupyter notebook provided in the Supporting Code). When fitting the bell-shaped model converged and returned plateau_2, logEC50_2, and dip within our constraints, we used the resulting values for the subsequent analysis.

If this bell-shaped model did not adequately match the data or failed to converge, the data were then fit to a conventional sigmoidal model (see Equation 6 below), as not all compounds tested induced a lower response at higher concentrations and were therefore unsuitable for a bell-shaped fit.

The minimum response estimated from the fitted curve was subtracted from all datapoints to calculate the net FRET signal for each compound and concentration. E_max_, defined as the maximum net FRET signal, and the corresponding pEC_50_ were then extracted from the fitted curve.

### BRET-based Go activation assay

We used the transiently transfected HEK-293T cell line for our BRET assays. Cells were grown in DMEM containing 10% fetal bovine serum, 2 mM L-glutamine, and 1% penicillin-streptomycin.

The BRET-based Gα_oA_ (Go) protein activation assay was performed as described previously.^43^ Briefly, Renilla luciferase 8 (Rluc8)-fused Gα_oA_ and mVenus-fused Gγ_2_ were used as the BRET pair and were cotransfected with Myc-tagged hMOR and G_β1_in the µg ratio 5:1:4:5 (hMOR:Gα_oA_:G_β1_:Gγ_2_). HEK293T cells were transiently transfected with the above constructs using polyethyleneimine (PEI) at a ratio of 2:1 (PEI:total DNA by weight). After ∼48 h of transfection, cells were washed, harvested, and resuspended in PBS + 0.1% glucose + 200 µM Na bisulfite buffer. 200,000 cells were then transferred to each well of the 96-well plates (White Lumitrac 200, Greiner bio-one) followed by addition of 1 µg/µL coelenterazine H, a luciferase substrate for BRET. Three minutes after addition of coelenterazine H, ligands were added to each well. Cells were incubated at 25^°^C within the PHERAstar *FSX* plate reader (BMG Labtech, Cary, NC, USA), with BRET signal measurements collected at 10 min. BRET ratio was calculated as the ratio of mVenus emission at 535 nm over RLuc8-catalyzed coelenterazine H emission at 475 nm. Data were collected from at least 4 independent experiments performed in triplicate.

Dose-response data were analyzed using a three-parameter sigmoidal dose response curve as follows:

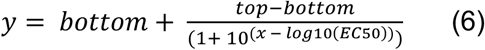

Net BRET ratios were calculated by subtracting the minimum response derived from the raw BRET curve. For each compound, the maximal net BRET response (E_max_) and corresponding pEC_50_ were extracted from the fitted curve. To compare efficacy across experiments, the E_max_ value for each compound was normalized to the E_max_ produced by DAMGO in the same experiment and expressed as %E_max_. %E_max_ and pEC_50_ values were then averaged across all independent experiments.

### Chemicals and reagents

Fentanyl, butonitazene, etonitazene, isotonitazene, N-desethyl isotonitazene, N-pyrrolidino etonitazene, 1-ethyl-pyrrolidinylmethyl N-desalkyl etonitazene, buprenorphine, and morphine were generously provided by Michael Baumann (National Institute on Drug Abuse (NIDA)). Cyclopropylfentanyl and valerylfentanyl were purchased from Sigma-Aldrich (St. Louis, MO, USA). N-desethyl etonitazene, N-pyrrolidino isotonitazene, etodesnitazene, and isotodesnitazene, and forskolin were purchased from Cayman Chemical (Ann Arbor, MI, USA). DAMGO was purchased from Tocris Bioscience (Minneapolis, MN, USA). Fluornitrazene and N-desethyl fluornitrazene were generously provided by Michael Michaelides (NIDA).

### Ligand preparation for molecular modeling and simulations

The 3D models of FNZ and its analogues were prepared using Ligprep of Schrodinger (version 2025-2). The protonation state of each ligand was assigned using Epik in Schrodinger (version 2025-2) under pH 7.0 ± 2.0 condition, which protonates the R2 amine nitrogen within the nitazene scaffold to its positively charged state.

### Conformational search

Conformational search was performed using MacroModel (Schrodinger Suite, version 2025-2) in an implicit water environment, employing the Mixed Torsional/Low-Mode sampling method and OPLS4 force field.^44^ The search was carried out with maximum 5,000 iterations and a convergence threshold of 0.005. Extended torsion sampling was enabled. Maximum steps = 10,000, and steps per rotatable bond = 500. All structures within 15 kJ/mol of the lowest energy conformer were considered in the final results.

For each nitazene analogue, the conformational search yielded several lowest-energy conformers because of symmetric flipping of the moieties. For presentation, we selected the lowest-energy conformer whose overall relative orientation of the three moieties matched that of the cryo-EM FNZ pose. All interatomic distances measurements reported were collected from these selected structures.

To evaluate the conformational difference between the lowest-energy conformer obtained from conformational search and the conformation within the MOR binding pocket for each nitazene analogue (see Results), representative conformations from the MD simulations were selected through visual inspection of the equilibrated trajectories. These conformers were subjected to extensive energy minimization using MacroModel to ensure the convergence to local minima, using parameters identical to those employed in the conformational search to allow direct energetic comparability.

### Quantum mechanical calculations

Quantum-mechanical calculations were performed using Density Functional Theory (DFT)^45^ with the Poisson-Boltzmann equation solved using Finite element methods (PB-FE) for implicit solvent treatment using Jaguar (Schrodinger Suite, version 2025-2). Calculations were conducted at the Ultrafine accuracy level. Electrostatic potential (ESP) surfaces and ESP derived atomic charges were computed. The resulting QM derived electron density maps were visualized to assess intrinsic differences among selected ligands and to observe the electrostatics of ligand conformations obtained from the MD simulations. All structures shown in Figures 4 and S4 were aligned to the benzimidazole core of FNZ using Maestro’s Superimpose function.

### Molecular dynamics simulations

The human MOR (hMOR)–G_i_ complex was constructed by homology modeling using MODELLER (version 10.6), with the cryo-EM structure of MOR bound to FNZ and complexed with the G_i_ protein used as the primary template.^24, 46^ A total of 25 hMOR models were generated, and the model with the lowest Discrete Optimized Protein Energy (DOPE) score was selected for following studies.^47^

The selected hMOR model was then processed through the Protein Preparation Wizard of Schrodinger as described previously.^48^ In particular, residues Asp116^2.50^ and Asp166^3.49^ were protonated to their neutral forms, consistent with assumptions for the active state of rhodopsin-like GPCRs, and Asp342^7.57^ was additionally protonated based on our previous work.^48^ Histidine protonation states were predicted by PROPKA and kept the same in all complexes, as summarized in Table S2. Using an equilibrated hMOR–FNZ model as the starting structure, we generated the other hMOR–nitazene models by manually modifying the FNZ structure with the corresponding substituents using the 3D Builder in Maestro (Schrodinger, version 2025-2).

To build the MD simulation systems, the hMOR-G_i_ complex models were immersed in an explicit 1-palmitoyl-2-oleoyl-sn-glycero-3-phosphocholine lipid bilayer (POPC) using the Desmond System Builder (Schrodinger, version 2025-2). The simple point charge (SPC) water model was used to solvate the system, the net charge of the system was neutralized by Cl^−^ ions, and then 0.15 M NaCl was added. The process resulted in a system with a dimension of 108×127x149 Å^3^ and total number of atoms of ∼195,000. We used the OPLS4 force field, while the initial parameters for all nitazenes were further optimized by the force field builder (Schrodinger, version 2025-2).

Desmond (D. E. Shaw Research, New York, NY) was used for the MD simulations. Similar to our previous simulation protocols used for GPCRs,^49^ the system was initially minimized and equilibrated with restraints on the ligand heavy atoms and protein backbone atoms. Simulations were carried out in the NPγT ensemble, with constant temperature (310 K) maintained using Langevin dynamics. Pressure was maintained at 1 atm using the hybrid Nose-Hoover Langevin piston method applied to an anisotropic flexible periodic cell. In the production runs, all restraints on the hMOR were released; however, to retain the integrity of the G_i_ protein, while allowing adequate flexibility for interaction with the receptor, the heavy atoms of residues 46-55, 182-189, and 230-242 of Gα, and the entire Gβ and Gγ subunits, were restrained with a force constant of 1 kcal/mol/Å^2^.

## Supporting Information

- Supporting Results: Stereochemistry-dependent R2 orientation underlies reduced activity of EP-ENZ
- Supporting Tables and Figures: Table S1, Summary of MD simulations; Table S2, The histidine protonation states of the MOR models; Table S3, Potential energy of EP-ENZ stereoisomers; Figure S1, Chemical structures of all compounds included in this study; Figure S2, Graphical representation of bell-shaped dose-response curve; Figure S3, Comparing Pose A and Pose B of the FNZ R1 tail; Figure S4, Conformational search results for MD-simulated ligands; Figure S5, MD supplemental material; Figure S6, α-Carbon chirality modulates EP-ENZ engagement with MOR
- Supporting Code: GTP curve-fitting code and examples

## Declarations of Competing Interests

No potential conflict of interest was reported by the authors.

## Author Contributions

M.J.R. collected and processed the experimental and computational modeling data, conducted the analysis, and wrote the manuscript. L.C. collected and processed the experimental data. A.T. collected and processed the computational modeling data, contributed to computational analysis and manuscript writing. K.H.L. contributed to conformational analysis. L.S. conceptualized and supervised the project, acquired funding, and revised the manuscript. All authors have reviewed and approved the final version of the manuscript.

## Acknowledgements

This research was supported by the National Institute on Drug Abuse–Intramural Research Program at the National Institutes of Health (NIH), Z1A DA000606 (L.S.). The contributions of the NIH author(s) are considered Works of the United States Government. The findings and conclusions presented in this paper are those of the author(s) and do not necessarily reflect the views of the NIH or the U.S. Department of Health and Human Services. We thank Michael Michaelides and Georgios Skiniotis for providing early access to the coordinates of the cryo-EM structure of the hMOR–Gi–FNZ complex prior to its publication. We also thank Sergi Ferré and Ning Sheng Cai for providing the FLP-FRT-HEK cell line stably expressing hMOR used in our cAMP inhibition assays, as well as the hMOR plasmid construct used in or BRET Go activation assay. This work utilized the computational resources of the NIH HPC Biowulf cluster (http://hpc.nih.gov).

## Notes

### Competing Interest Statement

The authors have declared no competing interest.

