## Supporting Information for "Coordinated subpocket engagement underlies nitazene potency at the µ-opioid receptor"

[#]equal contribution

### SUPPORTING RESULTS

#### Stereochemistry-dependent R2 orientation underlies reduced activity of EP-ENZ

EP-ENZ was the least efficacious nitazene in the GTP binding assay and was consistently ranked among lower-potency nitazenes in all assays. These findings prompted us to investigate whether EP-ENZ adopts conformations distinct from those of other nitazenes that might correlate with this reduced activity.

Because the carbon adjacent to the amine ($\alpha$-carbon) in EP-ENZ is chiral and the amino nitrogen can be protonated in two distinct configurations, we considered all four resulting stereochemical forms in our conformational searches to assess the impact of these different configurations. Comparison of the potential energies of the lowest-energy conformer for each enantiomer showed that changing the chirality of the $\alpha$-carbon had the greater effect (Table S3). Thus, we proceeded to analyze the 1R, 2R and 1S, 2R isomers (hereafter referred to as R-EP-ENZ and S-EP-ENZ, respectively), whose lowest energy conformers have the highest and lowest potential energies, respectively, to better understand the role of the $\alpha$-carbon chirality both as a free ligand and in the MOR-bound state.

In the conformational search, R- and S-EP-ENZ differed from other nitazenes in that their protons were oriented outwards, away from the ligand scaffold, and lacked engagement in cation-π interactions (Figure S6A). Instead, the R2 pyrrolidine ring engaged the 2-benzyl substituent in a CH-π interaction, which is less stabilizing than the analogous cation-π interactions observed in other analogues. The lower predicted energy of the S isomer relative to the R isomer is likely due to the ability of its pyrrolidine ring and ethyl groups to remain within 4 Å of the 2-benzyl substituents, whereas the R isomer could not engage in this hydrophobic interaction as effectively.

However, analysis of the equilibrated MOR-ligand complexes from our MD simulations showed that, when bound to the MOR, the proton of the S isomer was oriented inwards towards the 2-benzyl substituent and R1 alkoxy tail, preventing the ligand R2 group from forming a hydrogen bond with Asp149$^{3.32}$ (Figure S6B). Consequently, the positive electrostatic potential of S-EP-ENZ at the TM3 interface is significantly weaker compared to that of R-EP-ENZ, indicating a weaker salt bridge interaction between Asp149$^{3.32}$ and the ligand's R2 amine (Figure S6C). This orientation was maintained throughout the simulation, as shown in Figure S6D, where the proton consistently pointed inwards and remained disengaged from the receptor, as reflected by the larger distance between the proton and Asp149$^{3.32}$ than between the ligand nitrogen and Asp149$^{3.32}$ for the S enantiomer (Figures S6D and S6E).

As the strength of the salt bridge interaction between the positively charged amine and Asp149$^{3.32}$ is a hallmark of MOR recognition across most of the opioid scaffolds, this altered orientation in S-EP-ENZ impaired its ability to maintain high-affinity binding MOR, consistent with the reduced activity of the racemic mixture of EP-ENZ observed *in vitro*. Because EP-ENZ still exhibited nanomolar $EC_{50}$ across all assays, we propose that the observed activity is primarily driven by binding of the R stereoisomer, which is able to form a sufficiently strong ionic interaction.

### SUPPORTING TABLES

**Table S1: Summary of MD Simulations.**

Simulated time (ns) is an aggregation of trajectories for the indicated ligand, with $n_{trajectories} \geq 4$ for each compound. All trajectories employed the OPLS4 forcefield (see Methods).

| Ligand | Total time (ns) |
|---:|---:|
| FNZ | 3551 |
| ENZ | 3791 |
| DFNZ | 3642 |
| INZ | 2618 |
| DENZ | 4023 |
| DINZ | 3394 |
| BNZ | 4180 |
| EP-ENZ (R) | 3569 |
| EP-ENZ (S) | 3364 |

**Table S2: The histidine protonation states of the MOR models.**

The protonation states in the MOR models were predicted with PROPKA[1] at pH 7.0, as previously described.[2]

| Chain | Residue | Protonation state |
|---|---|---|
| Gα | H188 | HIP |
| | H195 | HIE |
| | H213 | HIE |
| | H244 | HID |
| | H322 | HID |
| Gβ | H54 | HIE |
| | H62 | HIE |
| | H91 | HIE |
| | H142 | HID |
| | H183 | HID |
| | H225 | HIE |
| | H266 | HIE |
| | H311 | HID |
| Gγ | H44 | HID |
| hMOR | H173 | HID |
| | H225 | HIE |
| | H299 | HID |
| | H321 | HIE |

**Table S3: Potential energy of EP-ENZ stereoisomers**

Potential energy of lowest-energy conformers of EP-ENZ's stereoisomers resulting from MacroModel's conformational search (Schrodinger Suite, version 2025-2).

| Conformer | Potential energy (kJ/mol) |
|---:|---:|
| $C_R N_R$ | 104.0 |
| $C_R N_S$ | 94.3 |
| $C_S N_R$ | 76.0 |
| $C_S N_S$ | 76.9 |

"R" and "S" denote the stereochemistry of "C", the $\alpha$-carbon, and "N", the protonated amine bonded to the $\alpha$-carbon in EP-ENZ (see Table 1, Figure S1, and Figure S6 for EP-ENZ's activity and full chemical structure).

### SUPPORTING FIGURES AND FIGURE LEGENDS

**Figure S1: Chemical structures of all compounds included in this study.**

Nitazenes and fentanyls are grouped in rows by their R1 pharmacophores, with parent comparators beginning each row.

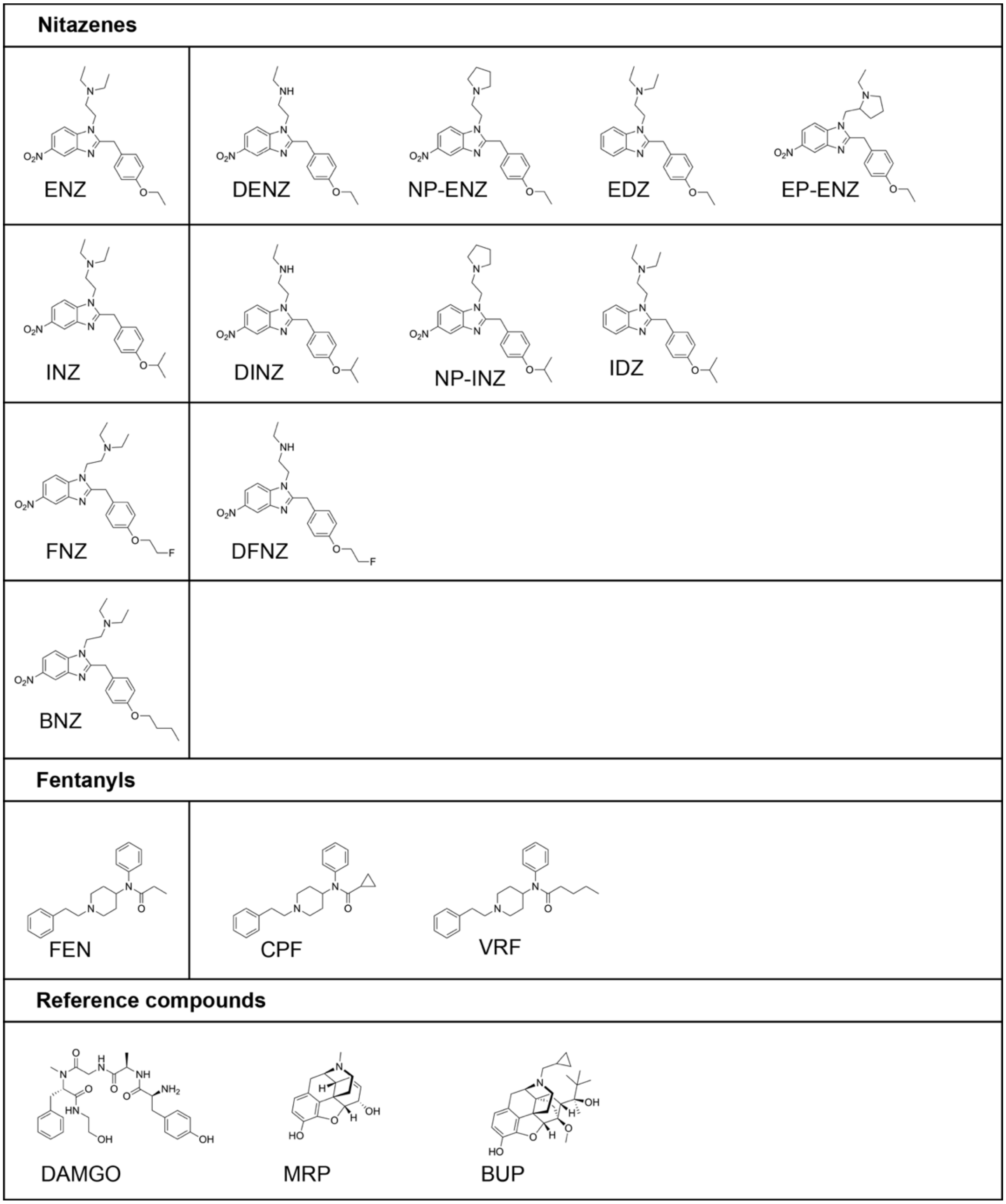

**Figure S2: Bell-shaped dose-response model applied to GTP-$G_i$ binding data.**

(A) Bell-shaped curve fit for individual, representative samples of opioids with varying efficacies tested in this study. INZ is a model bell-shaped (biphasic) response, whereas VRF exhibits a traditional sigmoidal dose-response. (B) Visual representation of the parameters employed in bell-shaped dose response regression equations (see Methods). Curve fit implementation can be found in the Supplementary Code.

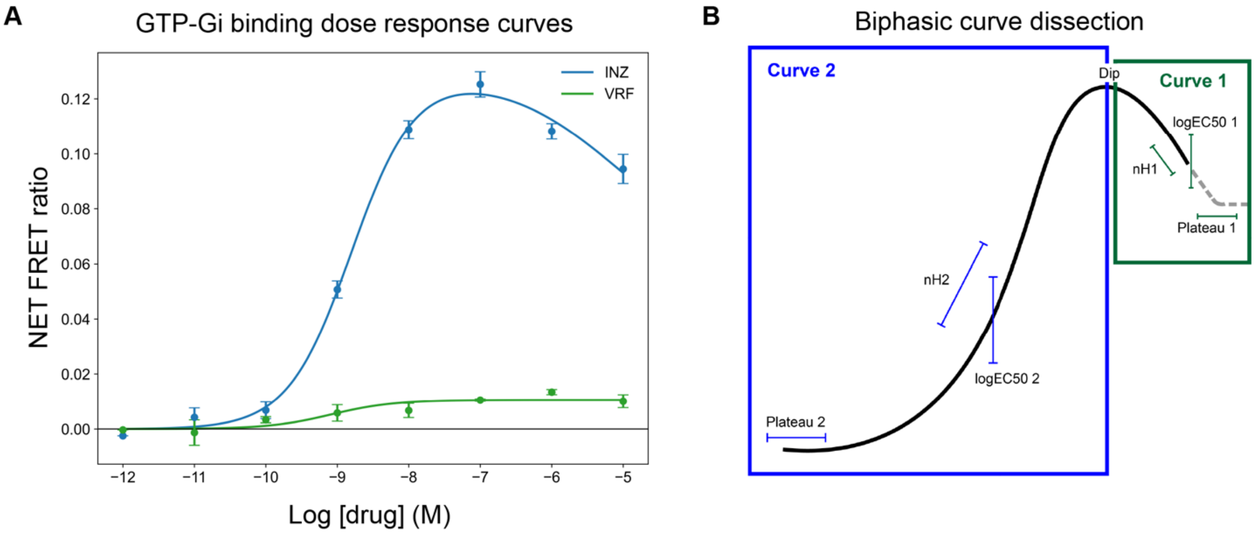

**Figure S3: Comparing Pose A and Pose B of the FNZ R1 tail.**

Depiction of the FNZ-MOR-$G_i$ cryo-EM structure as reported by Gomez and colleagues (PDB: 9O36).[3] We labeled the variation in R1 tail conformation as either Pose A (orange) or Pose B (purple) according to the PDB denotation and began our MD simulations from both poses.

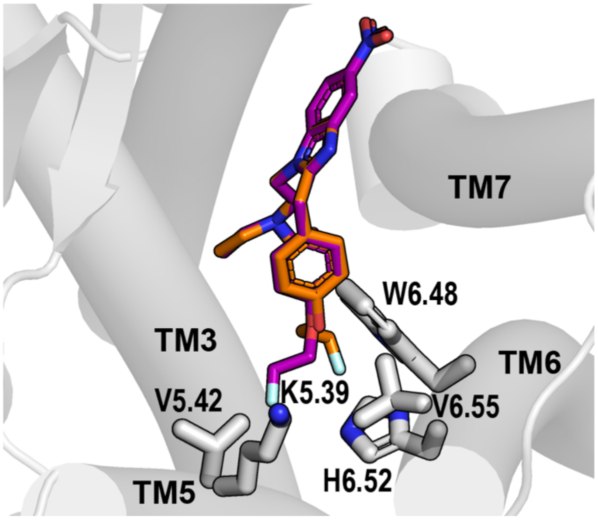

**Figure S4: Conformational search results for MD-simulated ligands.**

Yellow and pink highlights represent the R1 and R2 pharmacophores, respectively. R3 is defined as the 5-nitro group stemming from the benzimidazole core. The lowest-energy conformer of the indicated ligand is depicted, calculated from a molecular mechanics conformational search (see Methods). Proton orientation between N-diethyl and N-desethyl pairs varies systematically, with the proton orienting itself towards the 2-benzyl substituent in N-desethyl compounds, and towards the benzimidazole core in N-diethyl compounds, excluding BNZ.

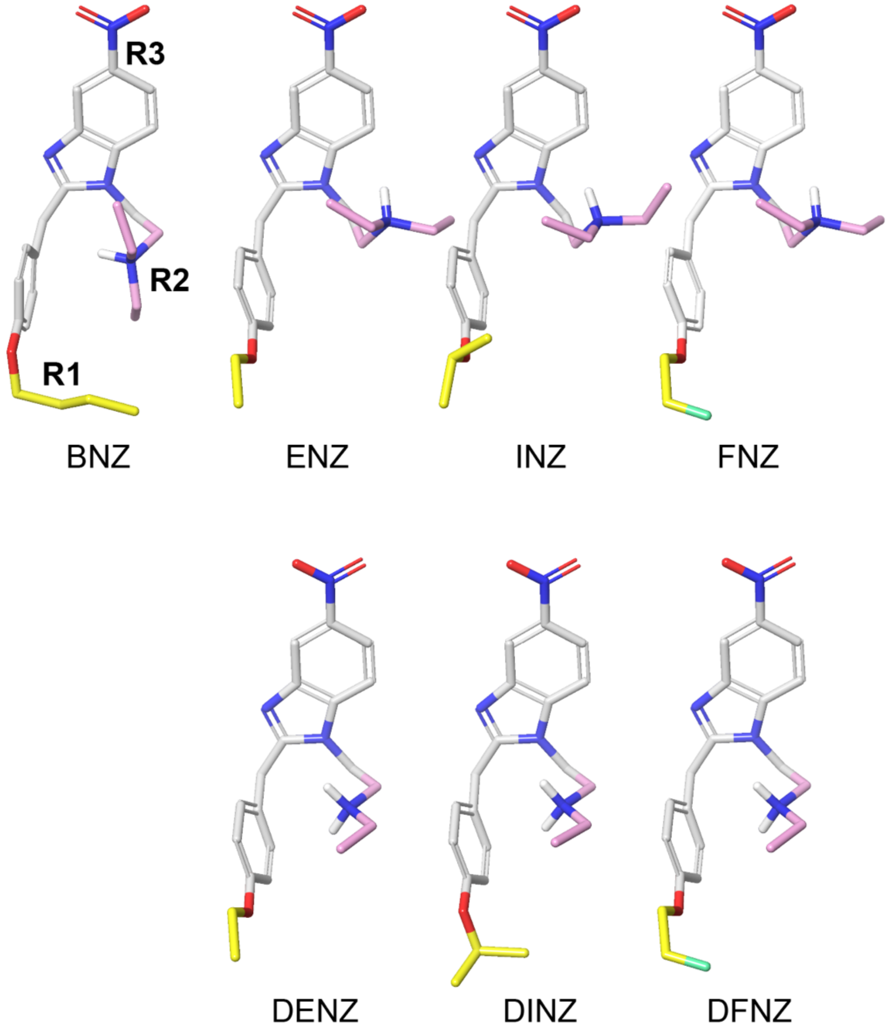

**Figure S5: R1-dependent effects of R2 N-desethylation in the ENZ/DENZ and FNZ/DFNZ pairs.**

Panels A-C compare the ENZ/DENZ pair, and panels D-F compare the FNZ/DFNZ pair, to show that the effects of R2 N-desethylation depend on the R1 substituent. (A,C) Regional contact frequencies and contact differences for DENZ versus ENZ (A) and DFNZ versus FNZ (C). (B, E) The contour plots of the distance distributions from each ligand's R2 protonated amine to the $C_\gamma$ of Asp149$^{3.32}$ and the OH of Tyr150$^{3.33}$. In both pairs, the N-desethyl analogue binds closer to Tyr150$^{3.33}$, with a larger shift for DFNZ relative to FNZ (E) than for DENZ relative to ENZ (B). Compared to DINZ (Figure 5D), all ligands depicted in this figure exhibit wider distance distributions, consistent with more dynamic engagement of the R2 subpocket. (C,F) Representative poses of ENZ and DENZ (C) and FNZ and DFNZ (F) from equilibrated trajectories, focusing on the R1 position. The R1 tails of DENZ (white and salmon) and ENZ (gray and fuchsia) occupy largely overlapping spaces within the R1 subpocket despite their differences at the R2 position. In contrast, the R1 tails of FNZ (gray and dark green) and DFNZ (white and light green) show slightly different spatial distributions within the R1 subpocket. However, compared to the INZ/DINZ pair (Figure 5F), the difference is less pronounced.

**A**

| | DENZ | | | | ENZ | | | | DENZ–ENZ | | | |
|---|---|---|---|---|---|---|---|---|---|---|---|---|
| | R1 | R2 | R3 | core | R1 | R2 | R3 | core | ΔR1 | ΔR2 | ΔR3 | Δcore |
| Y1.39 | | | 1.00 | 0.92 | | | 1.00 | 0.99 | | | | -0.07 |
| L2.57 | | | 0.98 | 0.83 | | | 0.94 | 0.91 | | | 0.04 | -0.08 |
| Q2.60 | | 0.37 | 0.98 | 1.00 | | 0.25 | 0.94 | 1.00 | | 0.12 | 0.04 | |
| S2.61 | | | 1.00 | 0.56 | | | 0.99 | 0.60 | | | 0.01 | -0.04 |
| N2.63 | | 0.02 | | | | | 0.14 | 0.14 | | 0.02 | -0.14 | -0.14 |
| Y2.64 | | 0.19 | 1.00 | 0.97 | | | 0.99 | 0.68 | | 0.19 | 0.01 | 0.29 |
| L2.65 | | | 0.17 | | | | 0.14 | | | | 0.03 | |
| D3.32 | | 0.98 | | 0.82 | | 1.00 | | 0.84 | | -0.02 | | -0.02 |
| Y3.33 | 0.10 | 0.93 | | 0.10 | 0.05 | 0.96 | | 0.02 | 0.05 | -0.03 | | 0.08 |
| M3.36 | 0.12 | 0.77 | | | 0.15 | 0.97 | | | -0.03 | -0.20 | | |
| K5.39 | 0.69 | | | | 0.57 | | | | 0.12 | | | |
| V5.42 | 0.88 | 0.18 | | | 0.94 | 0.40 | | | -0.06 | -0.22 | | |
| F5.43 | 0.36 | | | | 0.35 | | | | 0.01 | | | |
| W6.48 | 0.08 | 0.13 | | | 0.01 | 0.33 | | | 0.07 | -0.20 | | |
| I6.51 | 0.90 | 0.09 | | | 0.96 | 0.23 | | | -0.06 | -0.14 | | |
| H6.52 | 0.99 | | | | 0.99 | | | | | | | |
| V6.55 | 0.92 | | | | 0.97 | | | | -0.05 | | | |
| W7.35 | 0.42 | | | 0.48 | 0.48 | | | 0.40 | -0.06 | | | 0.08 |
| H7.36 | | | 0.97 | 0.70 | | | 0.99 | 0.87 | | | -0.02 | -0.17 |
| I7.39 | | 0.39 | 0.67 | 1.00 | | 0.80 | 0.49 | 1.00 | | -0.41 | 0.18 | |
| Y7.43 | | 0.84 | 0.32 | 0.99 | | 0.97 | 0.09 | 1.00 | | -0.13 | 0.23 | -0.01 |

**B**

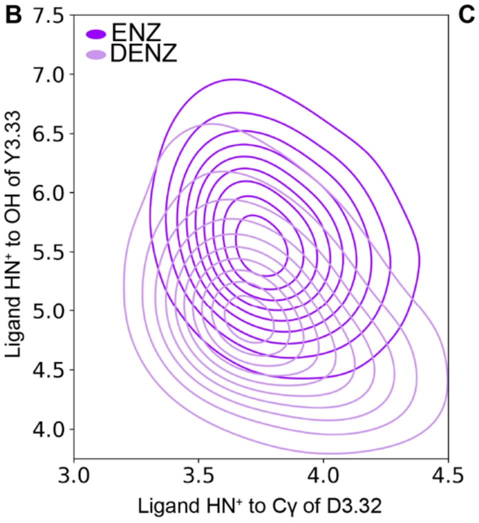


**C**

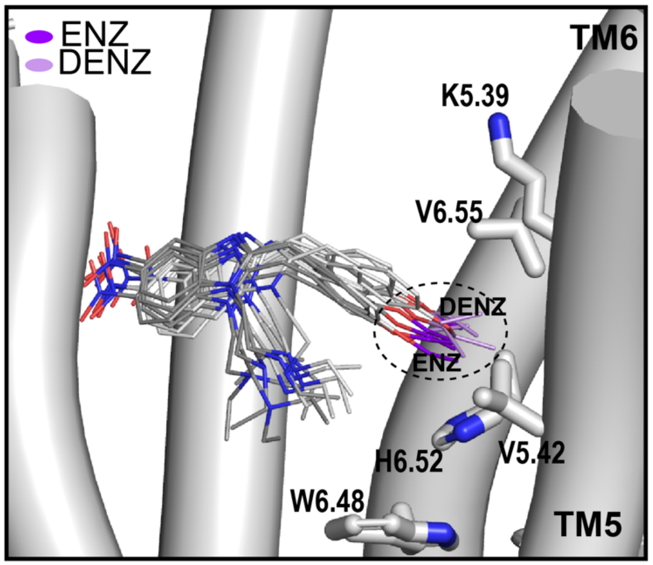


**D**

| | DFNZ | | | | FNZ | | | | DFNZ–FNZ | | | |
|---|---|---|---|---|---|---|---|---|---|---|---|---|
| | R1 | R2 | R3 | core | R1 | R2 | R3 | core | ΔR1 | ΔR2 | ΔR3 | Δcore |
| Y1.39 | | | 1.00 | 0.95 | | | 1.00 | 0.99 | | | | -0.04 |
| L2.57 | | | 0.98 | 0.90 | | | 0.97 | 0.97 | | | 0.01 | -0.07 |
| Q2.60 | | 0.27 | 0.92 | 1.00 | | 0.48 | 0.90 | 1.00 | | -0.21 | 0.02 | |
| S2.61 | | | 1.00 | 0.63 | | | 1.00 | 0.76 | | | | -0.13 |
| N2.63 | | | | | | | 0.50 | 0.49 | | | -0.50 | -0.49 |
| Y2.64 | | 0.04 | 1.00 | 1.00 | | 0.01 | 0.99 | 0.41 | | 0.03 | 0.01 | 0.59 |
| L2.65 | | | 0.39 | | | | 0.10 | | | | 0.29 | |
| D3.32 | | 1.00 | | 0.90 | | 1.00 | | 0.90 | | | | |
| Y3.33 | 0.22 | 0.80 | | 0.23 | 0.65 | 1.00 | | 0.01 | -0.43 | -0.20 | | 0.22 |
| M3.36 | 0.52 | 0.71 | | | 0.92 | 1.00 | | | -0.40 | -0.29 | | |
| K5.39 | 0.58 | | | | 0.02 | | | | 0.56 | | | |
| V5.42 | 0.96 | 0.21 | | 0.01 | 0.70 | 0.01 | | | 0.26 | 0.20 | | 0.01 |
| F5.43 | 0.37 | | | | 0.11 | | | | 0.26 | | | |
| W6.48 | 0.56 | 0.21 | | | 0.33 | 0.53 | | | 0.23 | -0.32 | | |
| I6.51 | 0.89 | 0.02 | | | 0.98 | 0.31 | | | -0.09 | -0.29 | | |
| H6.52 | 0.91 | | | | 0.99 | | | | -0.08 | | | |
| V6.55 | 0.77 | | | | 0.93 | | | | -0.16 | | | |
| W7.35 | 0.20 | | | 0.39 | 0.32 | | | 0.48 | -0.12 | | | -0.09 |
| H7.36 | | | 0.94 | 0.77 | | | 1.00 | 0.97 | | | -0.06 | -0.20 |
| I7.39 | 0.05 | 0.46 | 0.46 | 1.00 | 0.01 | 0.92 | 0.32 | 1.00 | 0.04 | -0.46 | 0.14 | |
| Y7.43 | | 0.97 | 0.28 | 1.00 | | 1.00 | 0.04 | 1.00 | | -0.03 | 0.24 | |

**E**

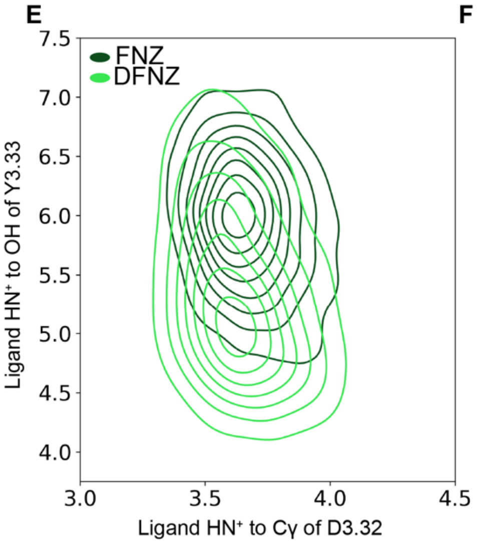


**F**

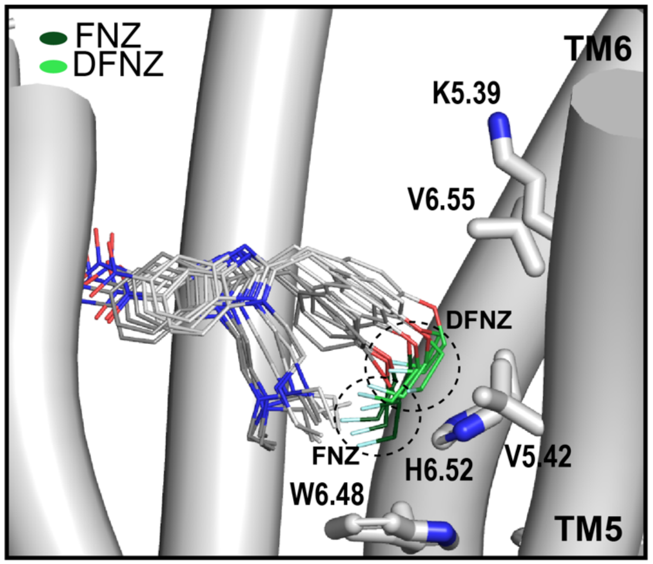

**Figure S6: α-Carbon chirality modulates EP-ENZ engagement with MOR.**

(A) Lowest energy conformers of 1S, 2R EP-ENZ (S-EP-ENZ) and 1R, 2R EP-ENZ (R-EP-ENZ) generated from MacroModel's conformational search (see Methods). "S" and "R" denote the chirality of the ligand's α-carbon S and R stereoisomers, respectively. (B) MD representative poses of S-EP-ENZ and R-EP-ENZ extracted from equilibrated simulations. (C) Electrostatic surfaces of the MD representative poses of S-EP-ENZ and R-EP-ENZ (note the molecules are in the same orientation as (B)). (D) Contour plots of the distance distributions from the proton (gray) and the nitrogen (blue) of the R2 amine to the Cγ of Asp149$^{3.32}$ for S- and R-EP-ENZ. In S-EP-ENZ, the nitrogen is closer to Asp149$^{3.32}$ than the proton, indicating that the ligand's proton is oriented inwards and is unable to form a hydrogen bond with the OH group of Asp149$^{3.32}$, although the salt bridge between the R2 amine and Asp149$^{3.32}$ remains. (E) Molecular graphic of S-EP-ENZ (pink proton) and R-EP-ENZ (yellow proton) as they interact with Asp149$^{3.32}$. Dotted line represents the hydrogen binding interaction between R-EP-ENZ's proton and the OH group of Asp149$^{3.32}$ graphed in panel (D).

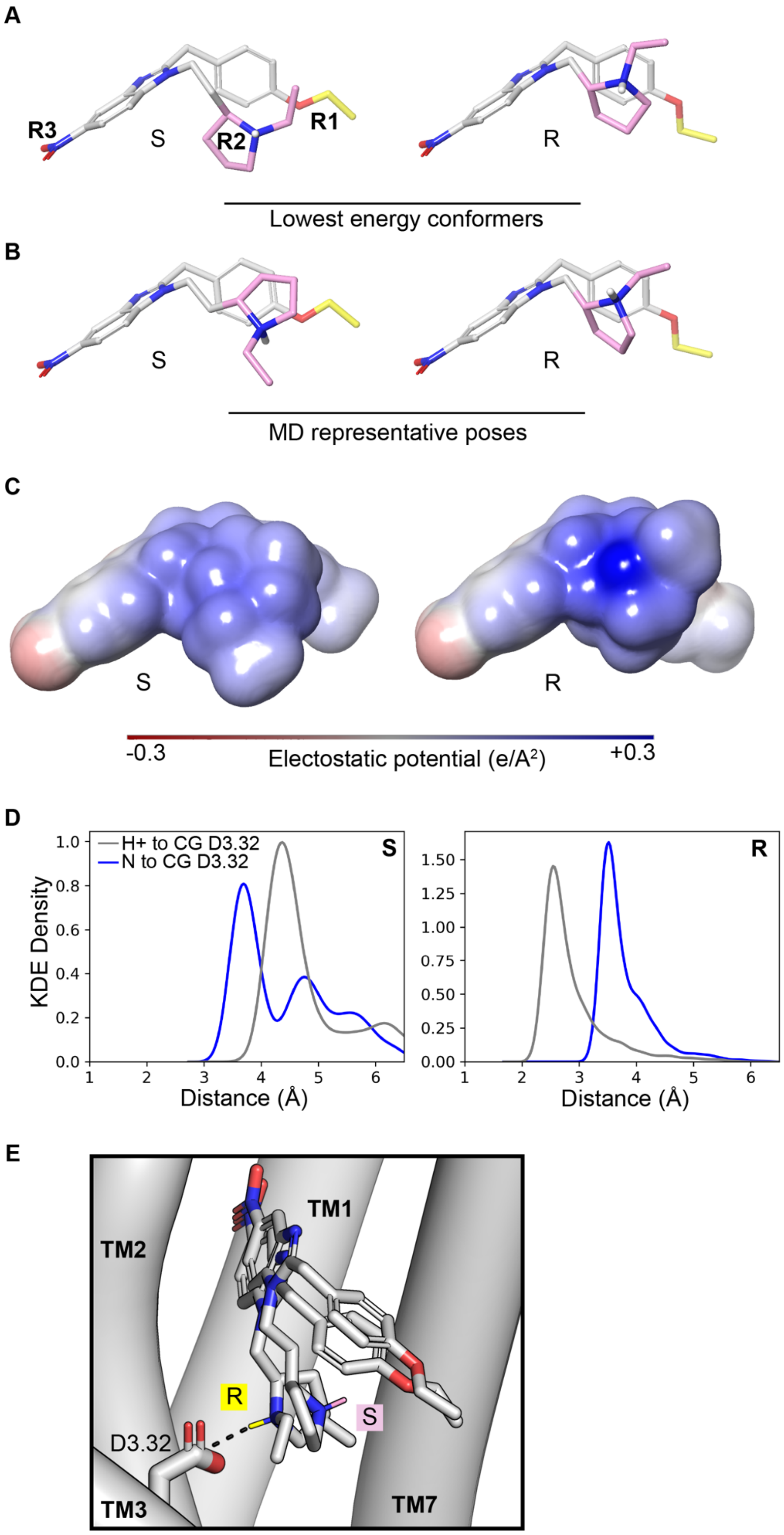
A
R3
S
R2
R1
R
Lowest energy conformers
B
S
R
MD representative poses
C
S
R
-0.3
Electostatic potential (e/A²)
+0.3
D
H+ to CG D3.32
N to CG D3.32
S
R
KDE Density
Distance (Å)
Distance (Å)
E
TM1
TM2
R
S
D3.32
TM3
TM7

### SUPPORTING CODE

#### GTP curve-fitting code and examples.

A Jupyter notebook, provided as a PDF export, implements the bell-shaped dose-response fitting procedure used in the analysis of the HTRF GTP-Gi binding data. The notebook includes example input data and representative fit outputs corresponding to the model described in the Methods and illustrated in Figure S2.
