## Supporting Code for "Coordinated subpocket engagement underlies nitazene potency at the µ-opioid receptor"

### GTP-Gi binding bell-shaped curve fit

This notebook contains the construction and implementation of our curve fit model adapted from GraphPad Prism's bell-shaped dose response curve. The data include two curves with varying efficacies and data shapes. The first curve, isotonitazene (INZ), is a model representation of the biphasic, or bell-shaped, response observed in this assay and is among our most efficacious compounds. The second curve, valerylfentanyl (VRF), is a known partial agonist and followed a traditional sigmoidal response, as demonstrated in the following graph. Running this notebook will produce a visual representation of dose-response data for both compounds.

##### Dependencies

This notebook requires the following Python libraries:

- `numpy`: For numerical operations and array handling.
- `pandas`: For data manipulation (though currently data is defined as numpy arrays).
- `matplotlib`: For plotting and visualization.
- `scipy`: For curve fitting using `curve_fit` and `minimize`.
- `lmfit`: For advanced model fitting with constraints.

You can install these dependencies using `pip`:

```
pip install numpy pandas matplotlib scipy lmfit
```

### Loading the necessary imports

In [1]:

```
import pandas as pd
import matplotlib.pyplot as plt
import numpy as np
from scipy.optimize import curve_fit, minimize
from lmfit import Model

def fit_bell(x, Plateau1, Plateau2, Dip, LogEC50_1, LogEC50_2, nH1, nH2):
    """
    Mathematical model for a bell-shaped (biphasic) dose-response curve.

    Args:
        x (float or np.ndarray): Log[Concentration] of the compound.
        Plateau1 (float): Initial plateau value (at low concentrations).
        Plateau2 (float): Final plateau value (at high concentrations).
        Dip (float): The maximum/minimum value of the curve's peak/trough.
```

```
        LogEC50_1 (float): Log of the effective concentration for the first
phase (descending curve).
        LogEC50_2 (float): Log of the effective concentration for the second
phase (ascending curve).
        nH1 (float): Hill slope for the first phase.
        nH2 (float): Hill slope for the second phase.

    Returns:
        float or np.ndarray: Predicted response value(s) for the given
Log[Concentration].
    """
    Span1 = Plateau1 - Dip
    Span2 = Plateau2 - Dip
    Section1 = Span1 / (1 + 10 ** ((LogEC50_1 - x) * nH1))
    Section2 = Span2 / (1 + 10 ** ((x - LogEC50_2) * nH2))

    return Dip + Section1 + Section2

def fit_sigmoid(x, bottom, top, logec50):
    """
    Mathematical model for a sigmoidal dose-response curve (Three-parameter
logistic).

    Args:
        x (float or np.ndarray): Log[Concentration] of the compound.
        bottom (float): Plateau at low concentrations.
        top (float): Plateau at high concentrations.
        logec50 (float): Log of the effective concentration for half-maximal
response.

    Returns:
        float or np.ndarray: Predicted response value(s) for the given
Log[Concentration].
    """
    y = bottom + (top - bottom) / (1 + np.power(10, (x - logec50)))
    return (y)

def calculate_r2(y, y_fit):
    """
    Calculates the R-squared (Coefficient of Determination) for a model fit.

    Args:
        y (np.ndarray): Actual observed data values.
        y_fit (np.ndarray): Predicted values from the model.
```

```
    Returns:
        float: R-squared value.
    """
    rss = np.sum((y - y_fit) ** 2)
    tss = np.sum((y - np.mean(y)) ** 2)
    r2 = 1 - (rss / tss)
    return r2


def _curvefitr2(x, y):
    """
    Core fitting logic that attempts a bell-shaped fit and falls back to a
sigmoid fit.

    This function utilizes `lmfit` to perform a bell-shaped fit with specific
constraints
    designed for GTP-Gi binding assays. If the bell fit fails, it attempts a
standard sigmoidal
    fit using `scipy.optimize.curve_fit`.

    Args:
        x (np.ndarray): Log[Concentration] data points.
        y (np.ndarray): Observed response data points.

    Returns:
        tuple: A tuple containing fit parameters and results:
               (plateau1, dip, logec50_1, plateau2, logec50_2, nh1, nh2, x2,
yfit, yfit_refined, r2)
               - plateau1 to nh2: Model parameters.
               - x2: High-resolution x-axis for plotting.
               - yfit: Predicted values for input x.
               - yfit_refined: Predicted values for x2.
               - r2: R-squared value of the fit.
    """
    x2 = []
    yfit = []
    yfit_refined = []

    x = np.array(x, dtype=float)
    y = np.array(y, dtype=float)

    mask = ~(np.isnan(y) | np.isinf(y))

    x_masked = x[mask]
    y_masked = y[mask]
```

```
    # initializing variables
    plateau2 = 0
    dip = 0
    logec50_2 = 0
    plateau1 = 0
    logec50_1 = 0
    nh2 = 1
    nh1 = 1
    r2 = 0

    try:
        plateau1_init = np.median(y_masked)
        plateau2_init = np.min(y_masked)
        dip_init = np.max(y_masked)
        logec50_1_init = np.median(x_masked)
        logec50_2_init = np.median(x_masked)
        nh1_init = 1
        nh2_init = 1

        # Constrains dip to greater than median, plateau 2, and second
highest y value
        median_y = np.median(y_masked)
        sorted_y = np.sort(y_masked)
        if len(sorted_y) > 1:
            second_highest_y = sorted_y[-2]
        else:
            second_highest_y = np.max(y_masked)
        dip_min_constraint = max(second_highest_y, 2 * plateau2_init, 1.5 *
median_y, 1e-6)

        # Constrains plateau 2 to be less than the fourth lowest y point
(purpose is to force bottom)
        if len(sorted_y) >= 4:
            plateau_2_max = sorted_y[3]  # Zero-based index, so this is the
4th lowest value
        else:
            plateau_2_max = np.median(y)

        bell = Model(fit_bell)

        params = bell.make_params(
            Plateau1=plateau1_init,
            Plateau2=plateau2_init,
            Dip=dip_init,
```

```
            LogEC50_1=logec50_1_init,
            LogEC50_2=logec50_2_init,
            nH1=nh1_init,
            nH2=nh2_init
        )
        params['Plateau1'].set(min=0)
        params['Plateau2'].set(min=0, max=plateau_2_max)
        params['Dip'].set(min=dip_min_constraint, max=np.max(y_masked))
        params['nH1'].set(min=0.5, max=5)
        params['nH2'].set(min=0.5, max=5)
        params['LogEC50_1'].set(min=0, max=np.max(x_masked))
        params['LogEC50_2'].set(min=np.median(x_masked) * 1.2,
max=np.max(x_masked))
        result = bell.fit(y_masked, params, x=x_masked)

        plateau1 = result.params['Plateau1'].value
        dip = result.params['Dip'].value
        logec50_1 = result.params['LogEC50_1'].value
        nh1 = result.params['nH1'].value
        plateau2 = result.params['Plateau2'].value
        logec50_2 = result.params['LogEC50_2'].value
        nh2 = result.params['nH2'].value

        yfit = fit_bell(x_masked, plateau1, plateau2, dip, logec50_1,
logec50_2, nh1, nh2)

        x2 = np.linspace(np.min(x_masked), np.max(x_masked), 500)
        yfit_refined = fit_bell(
            x2,
            plateau1,
            plateau2,
            dip,
            logec50_1,
            logec50_2,
            nh1,
            nh2
        )

        r2 = calculate_r2(y_masked, yfit)
        print("Bell fit successful")


    except ValueError as e:
        print(f"Bell fit failed with error: {e}")
```

```
        try:
            # Attempt sigmoid fit as fallback
            coeffs_init = (np.min(y_masked), np.max(y_masked),
np.median(x_masked))
            coeffs, pcov = curve_fit(fit_sigmoid, x_masked, y_masked,
coeffs_init, method='lm', maxfev=2000)
            bottom = coeffs[0]
            top = coeffs[1]
            logec50 = coeffs[2]

            yfit = fit_sigmoid(x_masked, bottom, top, logec50)

            # Generate refined fit for plotting
            x2 = np.linspace(np.min(x_masked), np.max(x_masked), 500)
            yfit_refined = fit_sigmoid(x2, bottom, top, logec50)

            # Compute R² value
            r2 = calculate_r2(y_masked, yfit)
            print(f'Sigmoid curve fit successful.')
            plateau2 = bottom
            logec50_2 = logec50
            dip = top

        except Exception as e:
            print(f'Curve fit failed with error: {e}')
            print('Using default values. Bottom as 0, Top as 0, logEC50 as 0,
r2 = 0.')
            r2 = 0
            x2 = []
            yfit = []
            yfit_refined = []

            plateau2 = 0
            dip = 0
            logec50_2 = 0
            plateau1 = 0
            logec50_1 = 0
            nh2 = 1
            nh1 = 0

    return plateau1, dip, logec50_1, plateau2, logec50_2, nh1, nh2, x2, yfit,
yfit_refined, r2

def plot_dose_response(
    data_dict,          # dict: {compound_name: np.ndarray}
```

```
    concentration_dict,      # dict: {compound_name: concentrations}
    colors,             # dict: {compound_name: color}
):
    """
    Processes dose-response data, performs curve fitting, and plots the
results.

    This function expects data organized by compound. It performs baseline
subtraction
    (relative to the last entry in the provided array), calculates mean and
SEM,
    fits either a bell-shaped or sigmoidal curve, and generates a
publication-quality plot.

    Args:
        data_dict (dict): Keys are compound names, values are numpy arrays
where
                         columns are replicates and rows correspond to
concentrations.
        concentration_dict (dict): Keys are compound names, values are
lists/arrays of
                                  Log[Concentration] values matching the
array rows.
        colors (dict): Keys are compound names, values are color strings for
plotting.

    Returns:
        list of dict: A list of dictionaries containing the fit parameters
for each compound.
    """

    param_list = []

    # Global plotting style
    plt.rcParams.update({
        "font.family": "Arial",
        "font.size": 14,
        "axes.labelsize": 16,
        "axes.titlesize": 18,
        "xtick.labelsize": 14,
        "ytick.labelsize": 14,
        "lines.linewidth": 2,
        "figure.dpi": 600
    })
```

```
    fig, ax = plt.subplots(figsize=(8, 6))

    for compound_name, conc in concentration_dict.items():
        concentrations = np.array(conc)
        compound = np.asarray(data_dict[compound_name])

        # Baseline subtraction
        baseline = compound[-1, :]
        compound_zeroed = compound - baseline

        # Mean + SEM
        mean_response = np.mean(compound_zeroed, axis=1)
        n_reps = compound_zeroed.shape[1]
        sem_response = np.std(compound_zeroed, axis=1, ddof=1) /
np.sqrt(n_reps)

        # Graph fitted curves
        x_fit = np.tile(concentrations, n_reps)
        y_fit_data = compound_zeroed.T.flatten()

        plateau1, dip, logec50_1, plateau2, logec50_2, nh1, nh2, \
        x2, yfit, yfit_refined, r2 = _curvefitr2(x_fit, y_fit_data)

        # --- Store parameters ---
        param_list.append({
            "compound": compound_name,
            "plateau1": plateau1,
            "dip": dip,
            "logEC50_1": logec50_1,
            "plateau2": plateau2,
            "logEC50_2": logec50_2,
            "Hill1": nh1,
            "Hill2": nh2,
            "R2": r2
        })

        # --- Align bottom to zero ---
        fit_min = np.min(yfit_refined)
        mean_shifted = mean_response - fit_min
        yfit_shifted = yfit_refined - fit_min

        # --- Plot data ---
        ax.errorbar(
            concentrations,
            mean_shifted,
```

```
            yerr=sem_response,
            fmt='o',
            color=colors[compound_name],
            capsize=5,
            markersize=6,
            zorder=3
        )

        # --- Plot fit ---
        ax.plot(
            x2,
            yfit_shifted,
            color=colors[compound_name],
            linewidth=2,
            label=compound_name
        )

    # --- Formatting ---
    ax.axhline(0, color='black', linewidth=1)
    ax.set_xlabel("")
    ax.set_ylabel("")
    ax.set_title("")
    ax.legend(frameon=False)

    plt.tight_layout()
    plt.show()

    return param_list
```

### Loading data

In [2]:

```
INZ = np.array([
    [0.209198018, 0.204912606, 0.198145561],
    [0.219958081, 0.214590965, 0.218757832],
    [0.239144754, 0.234962874, 0.230294397],
    [0.208606716, 0.222737819, 0.22370142],
    [0.162123248, 0.160186934, 0.158398988],
    [0.113642528, 0.121046064, 0.11447383],
    [0.117259823, 0.110324628, 0.113787991],
    [0.103698752, 0.106826802, 0.110711111]
])
VRF = np.array([
    [0.155252327, 0.148677919, 0.149890994],
    [0.154193099, 0.15549054, 0.154155911],
```

```
    [0.151507931, 0.151359746, 0.152208032],
    [0.14342946, 0.146123119, 0.154197755],
    [0.146770266, 0.151383356, 0.142894292],
    [0.145724128, 0.141917503, 0.145940904],
    [0.142394708, 0.145127799, 0.132120677],
    [0.140142262, 0.140255082, 0.14216354]
])
```

In [3]:

```
results = {}
data_dict = {
    'INZ': INZ,
    'VRF': VRF
}

concentration_dict = {
    'INZ': [-5, -6, -7, -8, -9, -10, -11, -12],
    'VRF': [-5, -6, -7, -8, -9, -10, -11, -12]

}

colors = {
    'INZ': '#1F77B4',
    'VRF': '#2CA02C',

}
```

### Running the curve fit

In [4]:

```
params = plot_dose_response(
    data_dict,
    concentration_dict,
    colors
)
Bell fit successful
Bell fit failed with error: Parameter 'Plateau2' has min == max
Sigmoid curve fit successful.
```

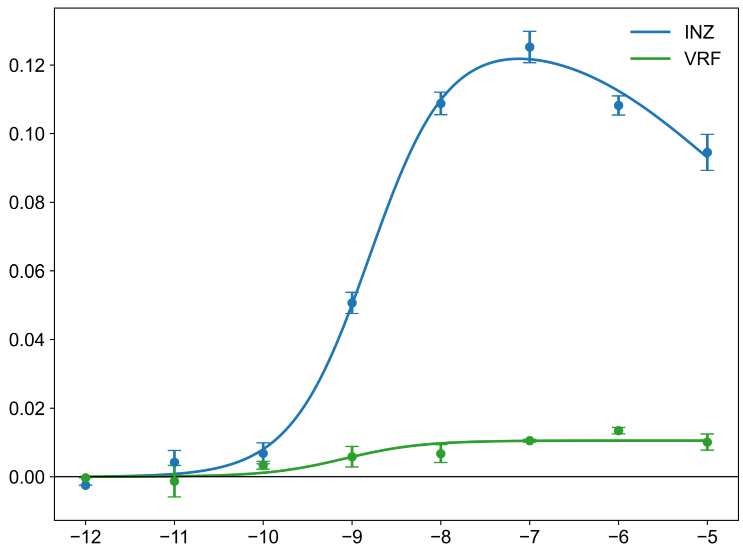

INZ
VRF
0.12
0.10
0.08
0.06
0.04
0.02
0.00
−12
−11
−10
−9
−8
−7
−6
−5
